# Only Zn^2+^ and Fe^2+^ out of 12 cations can fold ALS-linked nascent hSOD1

**DOI:** 10.1101/2023.04.29.538825

**Authors:** Liangzhong Lim, Jian Kang, Jianxing Song

**Affiliations:** Department of Biological Sciences, Faculty of Science, National University of Singapore, Republic of Singapore, 119260

**Author notes:** Correspondence to: Jianxing Song. L.Z.L. and J. K. contributed equally to this work. **Competing interests**: The authors declare no competing interests.

**Keywords:** Human CuZn-superoxide dismutase 1 (hSOD1), Oxidant stress, Amyotrophic lateral sclerosis (ALS), Cations, Protein folding and aggregation, NMR spectroscopy

## Abstract

153-residue copper-zinc superoxide dismutase 1 (hSOD1) is the first gene whose mutation was linked to FALS, while wild-type hSOD1 aggregation is associated with SALS. So far >180 ALS-causing mutations have been identified within hSOD1, but the underlying mechanism still remains enigmatic. Mature hSOD1 is extremely stable constrained by a disulfide bridge to adopt a Greek-key β-barrel fold that houses copper and zinc cofactors. Conversely, nascent hSOD1 is unfolded and prone to aggregation, requiring Zn^2+^ to initiate folding to a coexistence of folded and unfolded states. Recent studies demonstrate mutations disrupting Zn^2+^-binding correlate with their capacity to form toxic aggregates. Therefore, to decode the role of cations in hSOD1 folding provide not only mechanistic insights, but also therapeutic prospects for hSOD1-linked ALS. Here by NMR, we visualized the effect of 12 cations, encompassing 8 essential for humans (Na^+^, K^+^, Ca^2+^, Zn^2+^, Mg^2+^, Mn^2+,^ Cu^2+,^ Fe^2+^), 3 mimicking zinc (Ni^2+^, Cd^2+^, Co^2+^), and environmentally abundant Al^3+^. Surprisingly, most cations, including Zn^2+^-mimics, exhibited negligible binding or induction for folding of nascent hSOD1. Cu^2+^ displayed extensive binding to the unfolded state but induced severe aggregation. Unexpectedly, here for the first time Fe^2+^ was deciphered to have Zn^2+^-like folding-inducing ability. Surprisingly, Zn^2+^ failed to induce folding of H80S/D83S-hSOD1, while Fe^2+^ could. By contrast, Zn^2+^ could trigger folding of G93A-hSOD1, but Fe^2+^ failed. Notably, pre-existing Fe^2+^ disrupted the Zn^2+^-induced folding of G93A-hSOD1. Comparing with ATP-induced folded state, our results delineate that hSOD1 maturation requires: 1) intrinsic folding capacity encoded by the sequence; 2) specific Zn^2+^-coordination; 3) disulfide formation and Cu-load catalyzed by hCCS. This study decodes a previously-unknown interplay of cations in controlling the initial folding of hSOD1, thus not only underscoring the critical role of Zn^2+^ in hSOD1-associated ALS, but also suggesting novel hSOD1-dependent mechanisms for Cu^2+^/Fe^2+^-induced cytotoxicity likely relevant to other diseases and aging.

## Introduction

Amyotrophic lateral sclerosis (ALS) is the most prominent adult motor-neuron disease, clinically characteristic of progressive motor-neuron loss in the spinal cord, brainstem, and motor cortex, which leads to paralysis and death within a few years of onset. ALS was first described in 1869, which affects approximately 1-2 per 100,000 people worldwide (1–4). Most ALS cases are sporadic (90%) (SALS) whereas 10% are familial ALS (FALS). In 1993, CuZn-superoxide dismutase (SOD1) was identified to be the first gene associated with FALS (4), and its mutations cause the most prevalent form of FALS, accounting for ∼20% of total FALS cases (4–10). SOD1 is ubiquitously expressed in all tissues and is the most abundant protein in neurons comprising ∼1% of total protein. Currently, >180 mutations have been identified within the 153-residue human (hSOD1) that cause ALS by gain of toxicity (http://alsod.iop.kcl.ac.uk/) (9). Intriguingly, aggregation of the wild-type (WT) hSOD1 without any mutations has also been extensively found in SALS patients. So far, despite exhaustive studies, the exact mechanism for hSOD1-associated ALS still remains an enigma (4–14).

hSOD1 represents one of the most studied proteins not only for its physiological and pathological roles, but also for the fundamental principles of the enzymatic catalysis, protein folding and aggregation as well as their modulation by metalation (4–9). The mature hSOD1 is a homodimeric enzyme of remarkably high stability and solubility, with each subunit folding into an eight-stranded Greek-key β-barrel stabilized by an intramolecular disulfide bridge Cys57-Cys146. Each subunit holds one copper and one zinc ions in close proximity. While zinc ion is coordinated by His63, His71, His80, and Asp83, copper ion is ligated by His46, His48, and His120 (4–9) (Fig. 1A). Previous studies have established that nascent hSOD1 without metal cofactors and the disulfide bridge folds into the mature form through a very complex multi-step maturation process, whose detailed mechanism still remains to be completely elucidated. Nevertheless, it has been well recognized that the critical first step of the maturation is the initial folding of nascent hSOD1 specifically induced by Zn^2+^, which is followed by the formation of the disulfide bridge and incorporation of copper, both of which need the catalysis by copper chaperone for hSOD1 (CCS) (11–14). In contrast to mature hSOD1 of very high solubility, the early folding species particularly nascent hSOD1 have very high tendency of aggregation (4–16). Consequently, various super-stable hSOD1 mutants with Cys residues differentially mutated to Ala or/and Ser such as C6A/C111S mutant have been extensively used for biophysical and NMR structural studies (12).

**Fig. 1.**
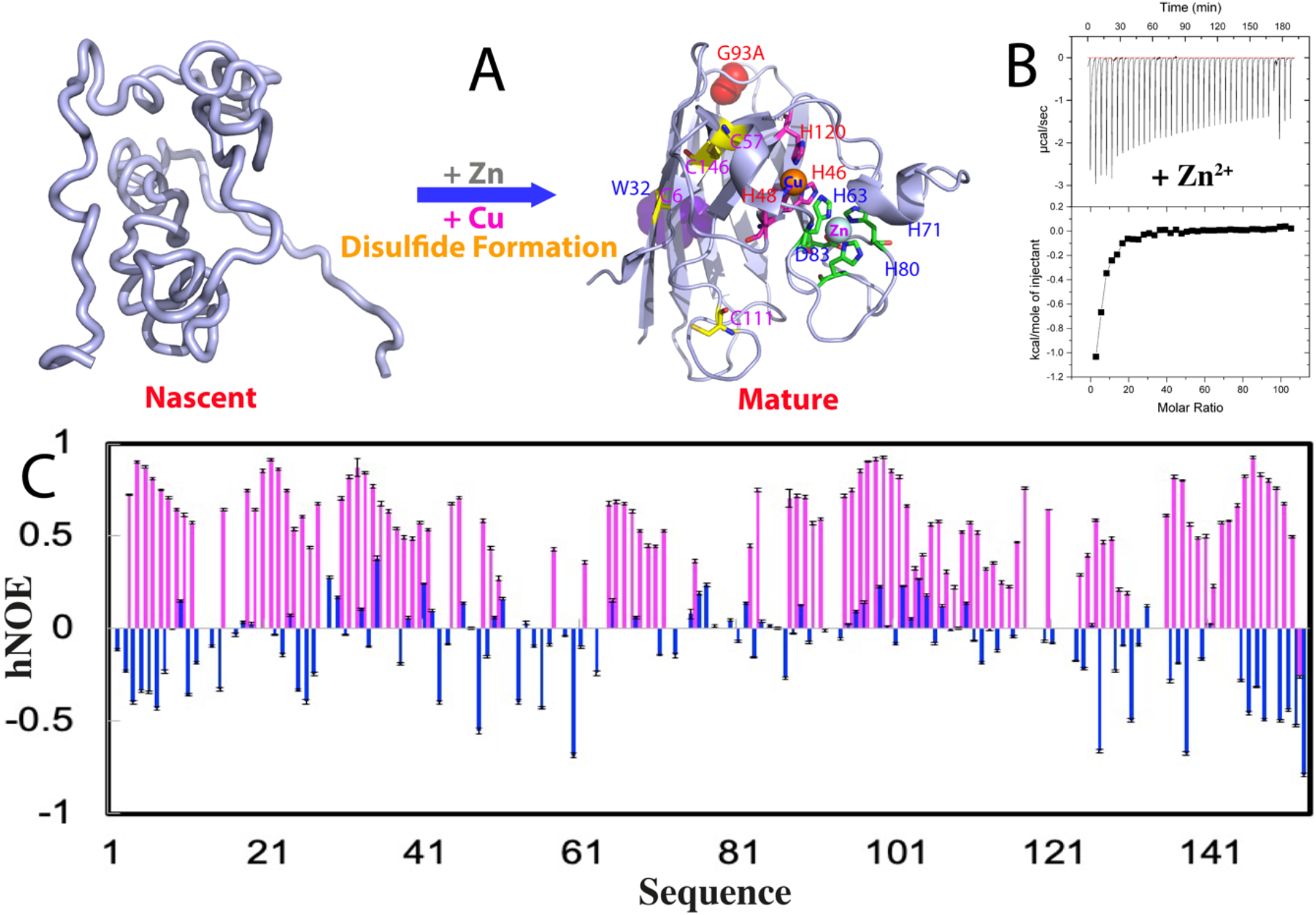
ITC and NMR characterization of the Zn^2+^-induced folding of nascent hSOD1. (A) Schematic representation of the folding of nascent hSOD1 into the folded subunit. The folded hSOD1 is represented by the monomer of the dimeric crystal structure of hSOD1 (1PU0). Residues for binding zinc and copper are labeled. (B) (Upper) ITC profiles of the interactions of the unfolded hSOD1 state with Zn^2+^, and (Lower) integrated values for reaction heats with subtraction of the corresponding blank results normalized against the amount of ligand injected versus the molar ratio of hSOD1: Zn^2+^. (C) {^1^H}–^15^N heteronuclear steady-state NOE (hNOE) of nascent hSOD1 (blue) and Zn^2+^-induced folded state (purple).

Previously, we have obtained atomic-resolution NMR evidence that L126Z-hSOD1, which has the C-terminal 28 residues truncated and considerably elevated toxicity compared to those with site substitution, was highly disordered without any stable secondary and tertiary structures, as well as lacking of restricted ps-ns backbone motions (15). We further found that nascent hSOD1 was also similarly disordered as L126Z-hSOD1 which is consistent with in-cell studies (11–14), thus providing a biophysical rationale for the observation that WT-hSOD1 is associated with SALS (16). Furthermore, we found that while nascent WT-hSOD1 could bind Zn^2+^ to undergo the initial folding into the co-existence between the folded and unfolded states at 1:20 (16), L126Z-hSOD1 completely lost the ability to interact with Zn^2+^ to fold (15). On the other hand, in the absence of Zn^2+^, nascent hSOD1 is also able to interact with membrane mimics including micelle and bicelle as L126Z-hSOD1. Nevertheless, upon availability of Zn^2+^, nascent hSOD1 becomes co-existing between two states and no longer able to interact with membrane mimics, while L126Z-hSOD1 remains disordered and still interacting with membrane mimics (15,16). Remarkably, *in vivo* studies revealed that the ALS-causing mutants of hSOD1 did initiate ALS pathogenesis by becoming associated with the ER and mitochondria membranes (17). Very recently, a biophysical study showed that the disruption of the ability of hSOD1 by ALS-causing mutations to bind Zn^2+^ is correlated to the acquired ability to become associated with mitochondria membrane to further form toxic aggregates (18). These results together highlight the extreme criticality of the initial folding of nascent hSOD1 induced by Zn^2+^, implying that the failure to achieve the initial folding renders WT-hSOD1 to become as toxic as the ALS-causing mutants, thus representing a convergent mechanism for the mutant hSOD1 to cause FALS and WT-hSOD1 to trigger SALS (15,16). On the other hand, accumulation of iron and copper within CNS was extensively identified in ALS cases (19–21). In particular, it has been revealed that in the SOD1-G93A mice, the breakdown of blood–spinal cord barrier (BSCB) in the early ALS disease phase led to accumulation of blood-derived iron in the spinal cord, which initiates ALS by triggering early motor-neuron degeneration through iron-induced oxidant stress (22). In this context, two questions of both fundamental and therapeutic importance arise: 1) in addition to Zn^2+^, are other cations also able to induce the initial folding of nascent hSOD1? 2) is hSOD1 involved in the manifestation of Fe2+-/Cu2+-cytotoxicity?

In this study, we aimed to address the two questions by NMR visualization. Briefly we have selected and evaluated the effects of 12 inorganic cations (Fig. S1), which include 8 (Na^+^, K^+^, Ca^2+^, Zn^2+^, Mg^2+^, Mn^2+^, Cu^2+^, and Fe^2+^) essential for humans (23), 3 (Ni^2+^, Cd^2+^ and Co^2+^) widely thought to have physicochemical properties very similar to Zn^2+^ and thus used to mimic Zn^2+^ in structure determination (5–8), as well as environmentally abundant Al^3+^. Surprisingly, most cations show no detectable ability to bind or to induce folding of nascent hSOD1 even up to 1:40 beyond which the sample started to precipitate. Intriguingly, Cu^2+^, the other cofactor of hSOD1, shows extensive binding to the unfolded state but triggers severe aggregation. Unexpectedly for the first time, Fe^2+^ has been identified to have the Zn^2+^-like ability to induce folding. Subsequently, we further characterized the effect of Fe^2+^ on the Zn^2+^-binding-defective H80S/D83S-hSOD1 as well as ALS-causing G93A-hSOD1 mutants. Very surprisingly, H80S/D83S-hSOD1 no longer undergoes the Zn^2+^-induced folding but can still be induced to fold by Fe^2+^. Conversely, Zn^2+^ retains its ability to initiate folding of G93A-hSOD1, whereas Fe^2+^ loses this capacity. Notably, the pre-existence of Fe^2+^ significantly disrupts the capacity of Zn^2+^ to induce the folding of G93A-hSOD1. These results together indicate that although the Zn^2+^- and Fe^2+^-binding pockets might be only partly overlapped, either binding is sufficient to induce folding of nascent hSOD1. Recently, we discovered that ATP and triphosphate could induce folding of nascent hSOD1, while ADP, Adenosine, pyrophosphate and phosphate, as well as TMAO had no detectable inducing activity (24). By comparing with the ATP-induced folded state of hSOD1, the results decipher that hSOD1 maturation requires: 1) intrinsic folding capacity encoded by the sequence; 2) additional information specifically provided by the Zn^2+^-coordination and; 3) disulfide formation and copper-load catalyzed by hCCS. Therefore, the current study not only unveils the irreplaceable role of Zn^2+^ in inducing the initial folding of nascent hSOD1 and preventing ALS pathogenesis, but also uncovers novel hSOD1-dependent mechanisms for Cu^2+^-/Fe^2+^-cytotoxicity.

## Results

### NMR characterization of the effect of Zn^2+^ on nascent hSOD1 and its H80S/D83S mutant

Nascent hSOD1 with the wild-type sequence is completely unfolded in the buffer, which has an HSQC spectrum with very narrow ^1^H and ^15^N spectral dispersions typical of a highly disordered protein (Fig. S2A). This is strikingly different from what was observed on various super-stable pseudo wild-type sequences in which four Cys residues (Cys 6, Cys 57, Cys 111 and Cys 146) were differentially replaced by Ala or Ser to increase stability and inhibit aggregation. These super-stable pseudo hSOD1 mutants including the C6A/C111S hSOD1 have been already folded to different degrees in the absence of Zn^2+^ ion (12).

As monitored by isothermal titration calorimetry (ITC), the heat release was observed upon adding Zn^2+^ (Fig. 1B), indicating that Zn^2+^ could interact with nascent hSOD1. However, as the heat changes are expected to result from at least two processes: namely binding of Zn^2+^ to hSOD1 and binding-induced folding, data fitting is not possible to obtain the thermodynamic parameters for the binding event. Consistent with ITC results, upon adding Zn^2+^, a folded population was formed as unambiguously indicated by the manifestation of a new set of well dispersed HSQC peaks (Fig. S2B). The formation of the folded state is largely saturated at a ratio of 1:20 (hSOD1: Zn^2+^) and even with further addition of Zn^2+^ up to 1:40, the unfolded and folded states still coexist (16), suggesting that Zn^2+^ alone is insufficient to completely convert the unfolded population into the folded state, consistent with the in-cell results that the complete formation of the mature WT-hSOD1 structure needs further disulfide formation and copper-load that is catalyzed by human copper chaperone for hSOD1 (hCCS) (10–14). Remarkably, upon adding EDTA at an equal molar concentration of Zn^2+^, the well-dispersed peaks of the folded state became completely disappeared, confirming that the formation of the folded state is the Zn^2^+-induced effect (16).

As shown in Fig. S2C, all residues of nascent hSOD1 have very small absolute values of (ΔCα–ΔCβ) chemical shifts, indicating the absence of any stable secondary structures (25). By contrast, many residues of the Zn^2+^-induced folded state have large and negative (ΔCα–ΔCβ) (Fig. S2C), which are highly similar to those of the Zn^2+^-induced state of the pseudo-WT-hSOD1 C6A/C111S except for several residues close to the mutation sites (Fig. S2D). The results together suggest that in the presence of Zn^2+^, both nascent WT and pseudo-WT hSOD1 adopt the highly similar β-barrel structures (Fig. S2E). Nevertheless, upon induction by Zn^2+^, all population of the pseudo-WT-hSOD1 became folded with a well-defined three-dimensional structure (Fig. S2E) as determined by NMR (12), which is very similar to the crystal structure of the mature hSOD1 (26). By contrast, WT-hSOD1 still has a co-existence of the unfolded and folded populations in the presence of an excess amount of Zn^2+^.

We further obtained {^1^H}–^15^N heteronuclear steady-state NOE (hNOE) of the folded and unfolded states of hSOD1 in the presence of Zn^2+^ at 1:20 (Fig. 1C), which reflects the backbone motion on ps-ns time scale (25,27–29). The residues of the unfolded state have small or even negative hNOE values with an average of -0.11, indicating that the unfolded state undergoes largely unrestricted backbone motions on ps-ns time scale. By contrast, most residues of the folded state have positive hNOE values with an average of 0.61, while some are even higher than 0.8, implying that the folded state has highly restricted backbone motions on ps-ns time scale. However, the average hNOE value of the folded state is much smaller than that of a well-folded protein such as EphA4 receptor (28), but similar to that of C71G-hPFN1 (24) and the N-terminal domain of the ALS-causing TDP-43 (29), which also have co-existing unfolded and folded conformations. Here, we further collected ^15^N-edited HSQC-NOESY spectrum on the sample with the coexistence of the unfolded and folded states of hSOD1 but unlike the C71G-hPFN1 (24) and N-terminal domain of TDP-43 (29), we found no NOE cross-peaks resulting from the exchange between two states, indicating that the time scale for the conformational exchange is slower than that for the C71G-hPFN1 (∼12 Hz) and TDP-43 N-Domain (∼14 Hz) (24,29,30).

We also generated H80S/D83S-hSOD1, which was previously shown to have severely-abolished capacity in Zn^2+^-binding (31). As shown in Fig. 2A, this mutant is also highly-disordered in the nascent form like the wild-type hSOD1, as evident by its poorly-dispersed HSQC spectrum with most peaks superimposable to those of the wild-type except for those of mutated residues and several residues close to the mutation sites in sequence. Indeed, upon addition of Zn^2+^ at a ratio of 1:10 (mutant: Zn^2+^), only very minor change was observed for the up-field 1D peaks (Fig. 2B) and the changes were mostly saturated at 1:30. However, even at a ratio of 1:40, the majority of well-dispersed HSQC peaks characteristic of the well-folded hSOD1 is not detected (Fig. 2C). Nevertheless, several relatively-dispersed HSQC peaks of the mutant in the presence of Zn^2+^ manifested and are mostly superimposable to those of the Zn^2+^-bound WT hSOD1 (Fig. 2D). The results together indicate that the mutations significantly reduce the zinc-binding capacity (31). Consequently, for this double mutant, even the excess supplement of Zn^2+^ could only trigger the formation of the partially-folded form.

**Fig. 2.**
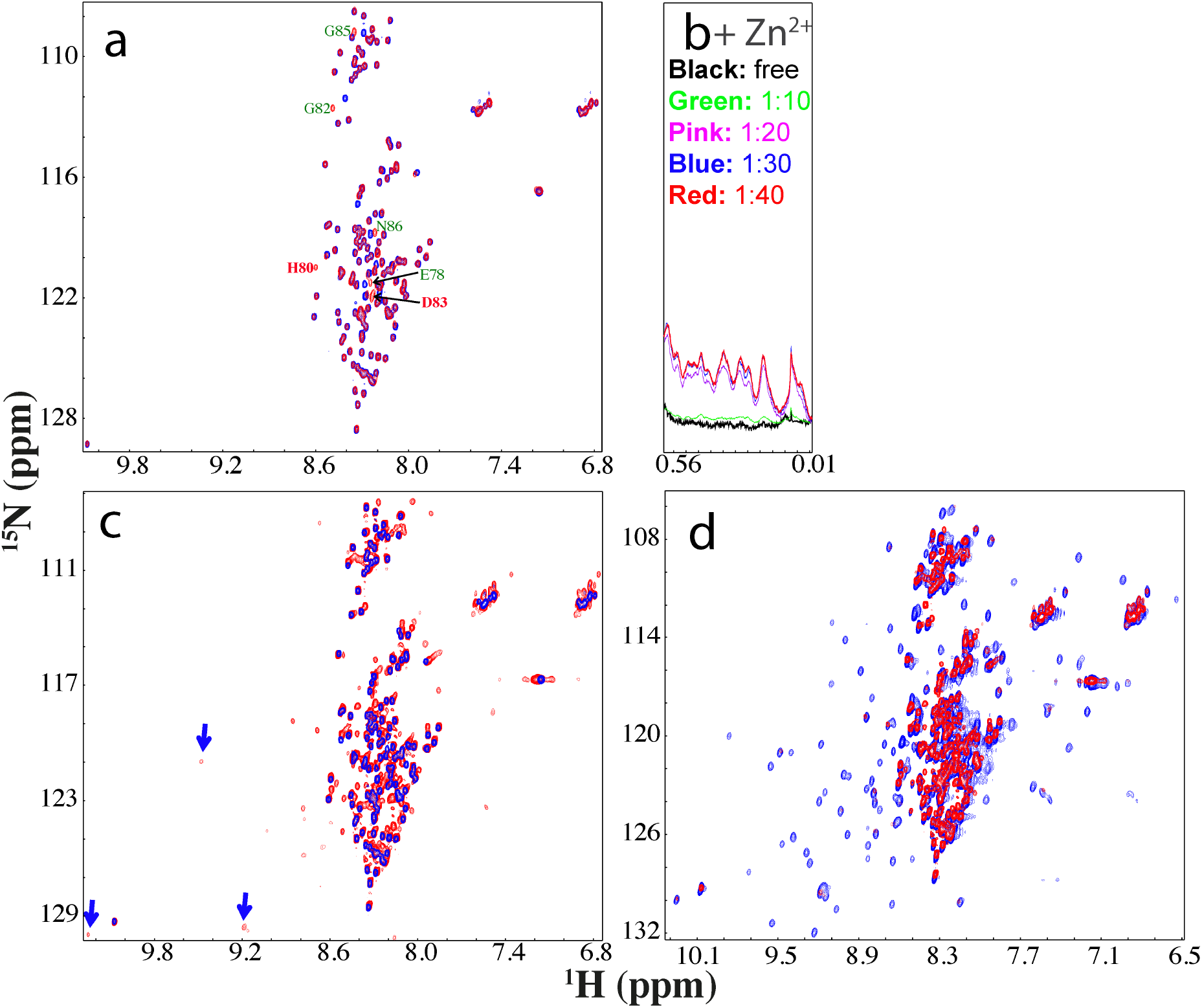
NMR characterization of the interaction of the H80S/D83S-hSOD1 with Zn^2+^. (A) Superimposition of HSQC spectra of the WT nascent hSOD1 (red) and H80S/D83S mutant (blue). Residues with large differences for their HSQC peaks are labeled. (B) Up-field 1D NMR peaks characteristic of the folded form of H80S/D83S (0.0-0.6 ppm) induced by the presence of Zn^2+^ at different molar ratios. (C) Superimposition of HSQC spectra of H80S/D83S in the absence (blue) and in the presence of Zn^2+^ at a molar ratio of 1:40 (red). Some well-dispersed peaks characteristic of the partially-folded form are indicated by blue arrows. (D) Superimposition of HSQC spectra of the WT hSOD1 in the presence of Zn^2+^ at a molar ratio of 1:20 (blue), and H80S/D83S in the presence of Zn^2+^ at a molar ratio of 1:40 (red).

### Copper extensively binds but triggers severe aggregation

We subsequently assessed whether copper, the other cofactor of hSOD1, can bind and induce folding of nascent hSOD1. Intriguingly, on the one hand, as shown in Fig. 3A, Cu^2+^ induced extensive shift and broadening of HSQC peaks of the unfolded state. On the other hand, however, unlike Zn^2+^ which is capable of inducing the manifestation of a complete set of well-dispersed HSQC peaks from the folded state, Cu^2+^ only induced the formation of a partially-folded state as indicated by the observations that only several well-dispersed HSQC peaks (Fig. 3A) and very up-field 1D peals (Fig. 3B) manifest such as that of Trp32 side chain which are from the folded state. The fact that most well-dispersed HSQC peaks characteristic of the Zn^2+^-induced folded state were undetectable implies that a large region of the Cu^2+^-induced state is not well-folded, or/and undergoes μs-ms conformational exchanges. Interestingly, the Cu^2+^-induced effect is mostly saturated at 1:6 (hSOD1:Cu^2+^) as evidenced by the very up-field 1D spectra. The current NMR results are generally consistent with the very recent report that unlike Zn^2+^, Cu^2+^ has no ability to prevent nascent hSOD1 from interacting with membrane to form toxic aggregates (18), implying that Cu^2+^ has no strong capacity to induce the initial folding of nascent hSOD1, which is required for abolishing the membrane-interacting ability.

**Fig. 3.**
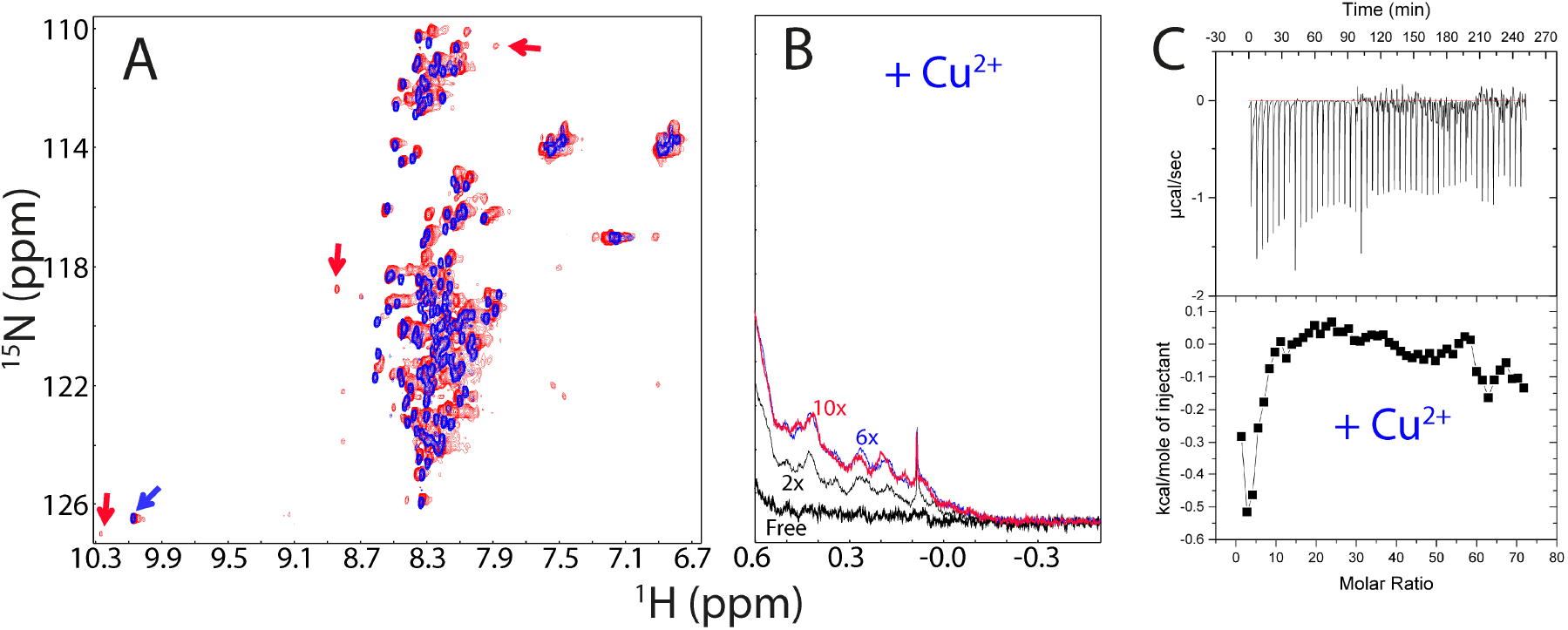
Cu^2+^ extensively binds the unfolded hSOD1 ensemble. (A) Superimposition of HSQC spectra of hSOD1 in the absence (blue) and in the presence of Cu^2+^ at a molar ratio of 1:10 (red). The blue arrow is used for indicating the HSQC peak of Trp32 ring proton characteristic of the unfolded ensemble and red ones for some HSQC peaks characteristic of the partially-folded form. (B) The up-field 1D NMR peaks characteristic of the partially-folded hSOD1 (-0.5-0.62 ppm) in the absence and in the presence of Cu^2+^ at different molar ratios. (C) (Upper) ITC profiles of the interactions of the unfolded hSOD1 state with Cu^2+^, and (Lower) integrated values for reaction heats with subtraction of the corresponding blank results normalized against the amount of ligand injected versus the molar ratio of hSOD1: Cu^2+^.

We also observed that even upon immediate addition of Cu^2+^, hSOD1 appeared to undergo μs-ms conformational exchanges or/and dynamic aggregation as evidenced by very broad 1D and HSQC peaks as compared with those in the presence of Zn^2+^ (Fig. 3A and 3B). Particularly after one hour, the NMR sample in the presence of Cu^2+^ started to form visible aggregates and consequently all NMR signals became disappeared. Consistent with this observation, addition of EDTA to the hSOD1 sample with freshly-added Cu^2+^ could result in the disappearance of very up-field 1D peaks and well-dispersed HSQC peaks, as well as the shifts of HSQC peaks, indicating that the observed effects are specifically induced by Cu^2+^. However, addition of EDTA to the Cu^2+^-added hSOD1 sample which already had visible aggregates failed to solubilize the aggregates as well as to restore the disappeared HSQC peaks.

We managed to conduct ITC titrations of Cu^2+^ into nascent hSOD1 sample at a low protein concentration (5 μM) and the result indicated that Cu^2+^ indeed showed the binding (Fig. 3C). Intriguingly, we also added Cu^2+^ to the hSOD1 sample in the pre-existence of Zn^2+^ at 1:20, but unfortunately the sample got precipitated immediately. In this context, the extensive binding of Cu^2+^ to the unfolded state, inability to induce the well-folded form, and strong induction of aggregation by Cu^2+^ may partly account for its high *in vivo* toxicity and also rationalize why the delivery and load of copper needs to be specifically implemented by hCCS to the pre-folded hSOD1 population (11–14).

### Most cations show no capacity in binding and inducing folding

So, a question of fundamental and biological interest is whether metal cations other than Zn^2+^ and Cu^2+^ can also bind and induce folding of nascent hSOD1. Previously, other metal cations have been extensively demonstrated to be capable of substituting either zinc or/and copper ions in mature hSOD1 (5–7) but it remains completely unknown whether they have the Zn^2+^-like capacity in triggering the folding of nascent hSOD1. To address this, we have further selected 10 other cations from the periodic table (Fig. S1A), which include Na^+^, K^+^, Ca^2+^, Mg^2+^, Mn^2+^, Fe^2+^, Ni^2+^, Cd^2+^, Co^2+^ and Al^3+^ and subsequently conducted titrations into nascent hSOD1 with ratios of cation:hSOD1 reaching up to 1:40 under the exactly same protein concentration and solution conditions used for Zn^2+^ as monitored by NMR HSQC spectroscopy, which can detect the binding events at residue-specific resolution with affinities ranging from very high to very low events even with Kd of mM (32,33).

Very unexpectedly, except for Fe^2+^, all cations triggered no considerable shift of HSQC peaks and no manifestation of very up-field 1D and well-dispersed HSQC peaks for nascent hSOD1, as exemplified by the results with Co^2+^, Ni^2+^, Cd^2+^, and Mn^2+^ which induce no large shift of HSQC peaks of nascent hSOD1 even with the ratio reaching up to 1:40 (Fig. 4), thus suggesting that these cations have no capacity to bind as well as to induce folding of nascent hSOD1. This is remarkably surprising because previously in various biophysical investigations and structure determinations, Ni^2+^, Cd^2+^ and Co^2+^ have been widely thought to have physicochemical properties very similar to Zn^2+^ and thus used to substitute Zn^2+^ in the mature hSOD1 (5–7). The present results thus strongly highlight the irreplaceable role of Zn^2+^ in initiating the first step of the maturation: folding of nascent hSOD1 (11–14). Indeed, previous folding studies have deciphered that zinc appears to uniquely modulate the entire folding free energy surface of hSOD1 (34).

**Fig. 4.**
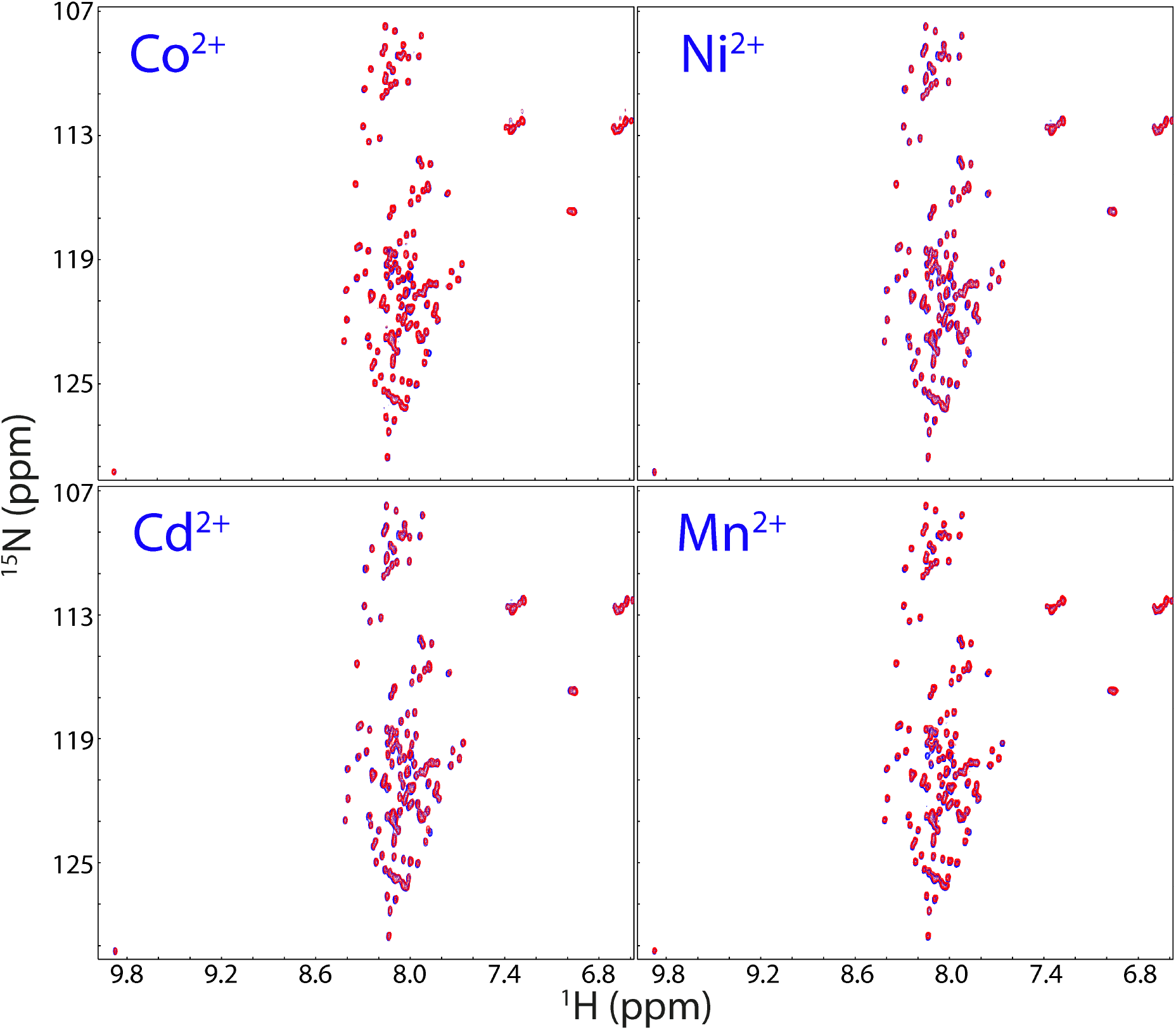
HSQC characterization of interactions of nascent hSOD1 with four cations. Superimposition of HSQC spectra of nascent hSOD1 (blue) and in the presence of Co^2+^, Ni^2+^, Cd^2+^, and Mn^2+^ respectively at a molar ratio of 1:40 (red).

So, an intriguing question arises: why does the WT hSOD1 still retain equilibrium between the folded and unfolded forms even in the excess presence of Zn^2+^? Previous studies revealed a surprising fact that on the one hand, to achieve copper-load and disulfide formation catalyzed by hCCS, the hSOD1 needs to populate the folded form bound to Zn^2+^ to some degree. Otherwise, the efficacy of maturation was low, as exemplified by the mutants (5–7,30,34,35). On the other hand, the interaction between hCCS and folded hSOD1 cannot be too stable but has to be dynamic/transit. If the SOD1-hCCS complex is too stable, the efficiency of the hCCS-catalyzed maturation will also be reduced or even abolished (36). So the request for dynamic/transit interactions between hSOD1 and hCCS appears to be elegantly fulfilled by the dynamic nature of the immature SOD1, which at least partly results from the co-existence of the unfolded and folded population in the presence of zinc, as we deciphered for the C71G-hPFN1 (24) and TDP-43 N-terminus (29).

### Fe^2+^ induces folding of nascent hSOD1

Here by a systematic assessment, for the first time we found that most unexpectedly Fe^2+^ owns the capacity in triggering folding of nascent hSOD1 into an equilibrium between the unfolded and Fe^2+^-induced folded states. So, we subsequently conducted a detailed NMR characterization to elucidate its conformational features. As seen in Fig. 5A, addition of Fe^2+^ triggered the manifestation of very up-field 1D and well-dispersed HSQC peaks characteristic of the folded form and the increase in the folded population is mostly saturated at a molar ratio of 1:20 (hSOD1: Fe^2+^). Fig. 5B presents the superimposition of the HSQC spectra of the Fe^2+^- and Zn^2+^-bound hSOD1 forms. Unfortunately, a detailed comparison of the intensities and chemical shifts of NMR peaks induced by Fe^2+^ and Zn^2+^ is impossible because Fe^2+^ has strong paramagnetic effects including effects of relaxation enhancement and pseudo-contact shift (37), which also made the sequential assignment of the Fe^2+^-bound state impossible by NMR. However, ITC measurement also revealed that Fe^2+^ could indeed bind hSOD1 (Fig. 5C).

**Fig. 5.**
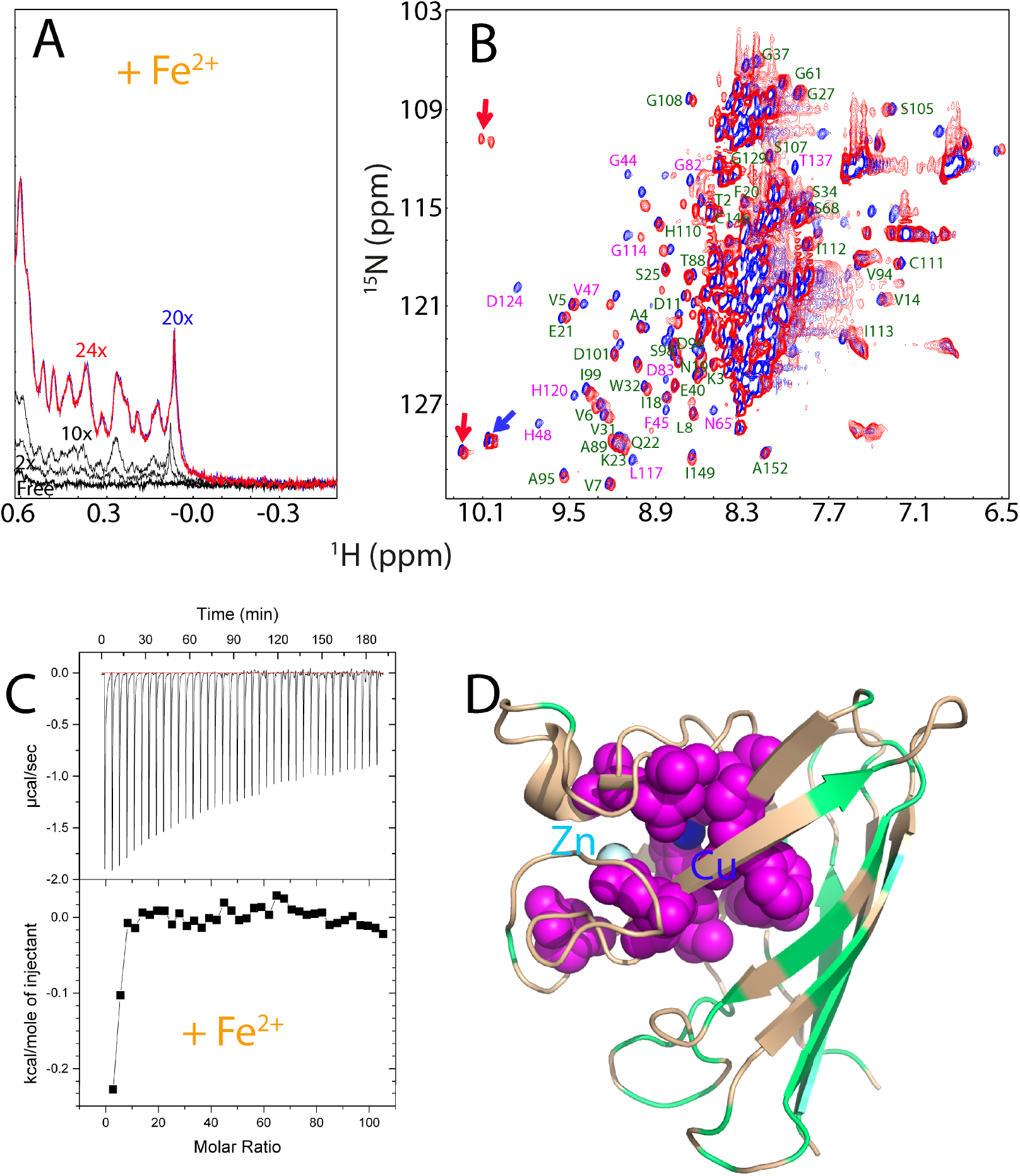
Fe^2+^ has the Zn^2+^-like capacity in triggering the folding of nascent hSOD1. (A) Up-field 1D NMR peaks characteristic of the folded hSOD1 (-0.5-0.62 ppm) in the absence and in the presence of Fe^2+^ at different molar ratios. (B) Superimposition of HSQC spectra of hSOD1 in the presence of Zn^2+^ (blue), and Fe^2+^ (red). The labels of the sequential assignments for some well-resolved HSQC peaks are in green if the peaks are largely superimposable but in pink if the peaks are largely shifted in both forms. (C) (Upper) ITC profiles of the interactions of the unfolded hSOD1 ensemble with Fe^2+^, and (Lower) integrated values for reaction heats with subtraction of the corresponding blank results normalized against the amount of ligand injected versus the molar ratio of hSOD1: Fe^2+^. (D) The monomer structure of hSOD1 (1PU0) with the residues displayed in pink spheres if their HSQC peaks showed large differences in the Zn^2+^- and Fe^2+^-forms, and colored in green if their HSQC peaks were superimposable.

Nevertheless, based on the assignments of the Zn^2+^-bound form, many peaks from the Fe^2+^-bound hSOD1 are highly superimposable to those of the Zn^2+^-induced state; while some have large differences. Interestingly, as shown in Fig. 5D, the peaks with significant differences are from the residues surrounding the Cu/Zn binding pocket, while the peaks without significant differences are from those located relatively far away from the Cu/Zn binding pocket such as over the β-barrel. This strongly implies that the Fe^2+^- and Zn^2+^-induced hSOD1 structures might be very similar particularly over the β-barrel regions, and further implies that Fe^2+^-binding pocket is located close to, or even has overlaps with the Cu/Zn binding pocket. Consequently, the residues close to Fe^2+^ showed large differences for their HSQC peaks due to the pseudo-contact shift effect (37), or/and slightly-different conformations in the Fe^2+^-induced state. Moreover, the up-field and well-dispersed HSQC peaks also became completely disappeared upon adding EDTA, confirming that the formation of the folded hSOD1 is also Fe^2+^-induced effect.

We further conducted a competitive experiment between Zn^2+^ and Fe^2+^ by monitoring the changes of both up-field (Fig. 6A) and HSQC (Fig. 6B and 6C) peaks upon stepwise addition of Zn^2+^ into the hSOD1 sample in the pre-existence of 20x Fe^2+^. As seen in Fig. 6A, with addition of 10x zinc, the zinc-specific peak in the 1D spectrum manifested. On the other hand, however, as shown in HSQC spectra (Fig. 6B), in the presence of 10x zinc, many HSQC peaks still remain highly similar to those of the Fe^2+^-induced state. Upon addition of 20x Zn^2+^, although many peaks become superimposable to those of Zn^2+^-bound form, there are still some peaks more superimposable to those of the Fe^2+^-induced state (Fig. 6C). This implies that Fe^2+^ and Zn^2+^ binding-sites might be only partly overlapped. Unfortunately, further addition of Zn^2+^ induced severe broadening of the NMR peaks and then visible aggregates of the sample, thus blocking further characterization. Nevertheless, the results of the competitive experiment strongly imply that the presence of Fe^2+^ at high concentrations can interfere in the Zn^2+^-induced formation of the well-folded hSOD1, which may reduce the efficiency of the maturation of WT hSOD1.

**Fig. 6.**
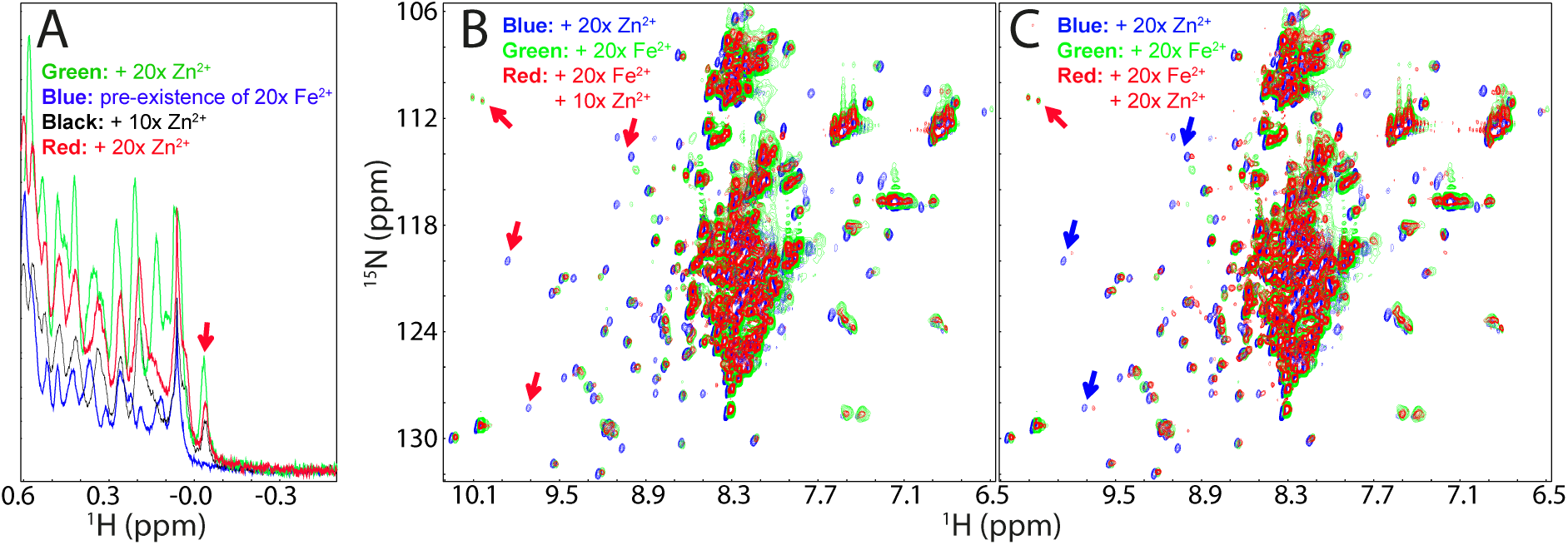
Fe^2+^ and Zn^2+^ have partly overlapped binding sites. (A) Up-field 1D NMR peaks (-0.5-0.62 ppm) of hSOD1 in the presence of 20x Zn^2+^ (green), 20x Fe^2+^ (blue), addition of 10x Zn^2+^ into the sample in the pre-existence of 20x Fe^2+^ (black); and addition of 20x Zn^2+^ into the sample in the pre-existence of 20x Fe^2+^ (red). (B) Superimposition of HSQC spectra of hSOD1 in the presence of 20x Zn^2+^ (blue), in the presence of 20x Fe^2+^ (green), and in the presence of both 20x Fe^2+^ and 10x Zn^2+^ (red). (C) Superimposition of HSQC spectra of hSOD1 in the presence of 20x Zn^2+^ (blue), in the presence of 20x Fe^2+^ (green), and in the presence of both 20x Fe^2+^ and 20x Zn^2+^ (red). Red arrows are used for indicating Fe^2+^-specific HSQC peaks while blue arrows for Zn^2+^-specific peaks.

Intriguingly, we also attempted to add Cu^2+^ into the hSOD sample in the pre-existence of 20x Fe^2+^. Unfortunately, even the addition of Cu^2+^ at 1:5 induced severe aggregation of the sample and NMR signal became undetectable.

### Characterization of the Zn^2+^- and Fe^2+^-binding pockets with hSOD1 mutants

We further titrated the H80S/D83S-hSOD1 mutant with Fe^2+^ by monitoring the changes of up-field 1D (Fig. 7A) and HSQC (Fig. 7B) spectra. Very unexpectedly, although for this mutant, Zn^2+^ failed to trigger the formation of the well-folded structure (Fig. 2), Fe^2+^ is still able to induce folding as evident from the manifestation of up-field peaks mostly saturated at a ratio of 1:20 (Fig. 7A), as well as well-dispersed HSQC peaks (Fig. 7B), which are mostly superimposable to those of WT-hSOD1 (Fig. 7C). These results suggest that the Fe^2+^-induced structures are highly similar for both H80S/D83S mutant and wild-type hSOD1. Crucially, the results also reveal that the residues for binding Zn^2+^ and Fe^2+^ are not identical. Very strikingly, however, either Zn^2+^- or Fe^2+^-binding is sufficient to trigger folding of nascent hSOD1.

**Fig. 7.**
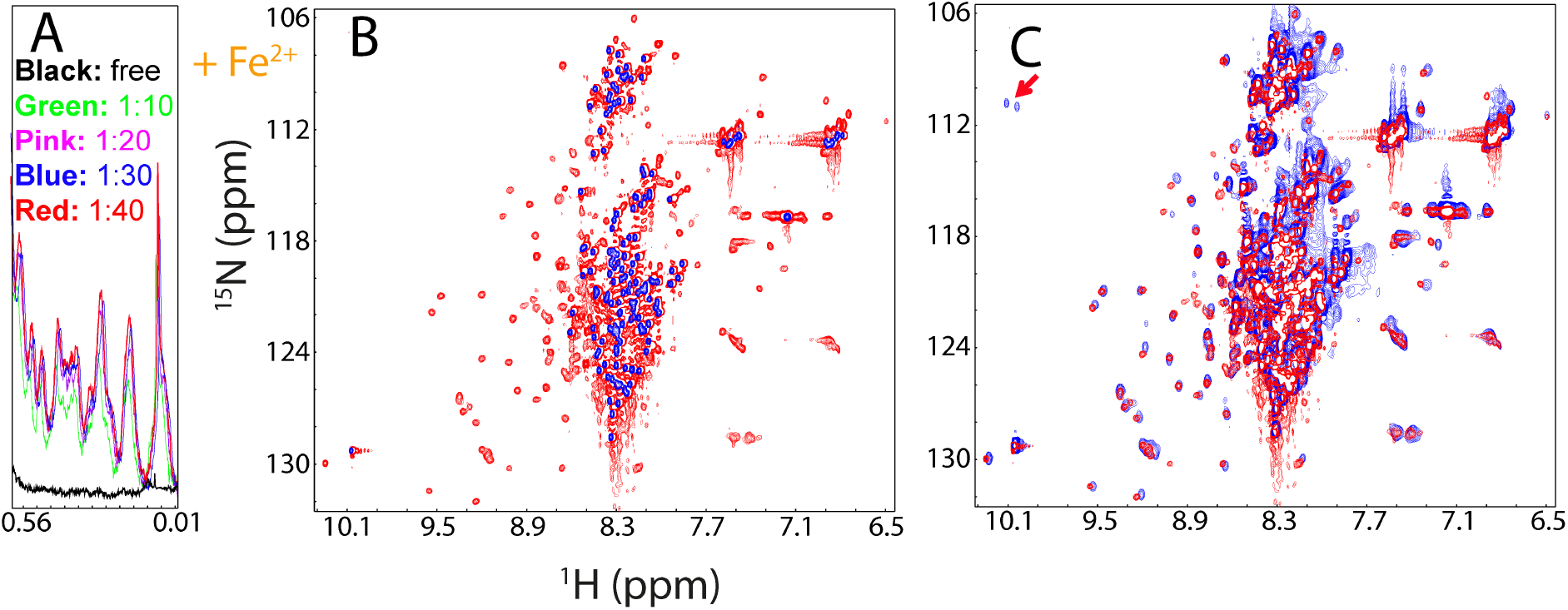
NMR characterization of the interaction of the H80S/D83S hSOD1 with Fe^2+^. (A) Up-field 1D NMR peaks characteristic of H80S/D83S hSOD1 (0.0-0.6 ppm) in the absence and in the presence of Fe^2+^ at different molar ratios. (B) Superimposition of HSQC spectra of H80S/D83S-hSOD1 in the absence (blue) and in the presence of Fe^2+^ at a molar ratio of 1:20 (red). (C) Superimposition of HSQC spectra of the WT hSOD1 (blue) and H80S/D83S-hSOD1 (red) in the presence of Fe^2+^ at a molar ratio of 1:20.

### Fe^2+^ radically disrupts the Zn^2+^-induced folding of G93A-hSOD1

So far, >180 ALS-causing mutations have been identified on the 153-residue hSOD1 but it still remains highly elusive for the mechanism by which these mutations cause ALS. To explore this, we generated the ALS-causing G93A-hSOD1 protein, which is also highly disordered like WT as judged by its poorly-dispersed HSQC spectrum (Fig. 8A). Interestingly, however, in addition to the residues such as Asp92, Val94 and Ala95 which are next to the mutation residue Gly93 in sequence, many residues such as Gln22, Glu24, Asp101 and Asp103 far away from Gly93, also have significant shifts for their HSQC peaks (Fig. 8A). This is very different from what observed on the double mutant H80S/D83S (Fig. 2a), implying that even in the unfolded ensemble, the ALS-causing mutation of G93A is able to trigger long-range perturbations on other residues.

**Fig. 8.**
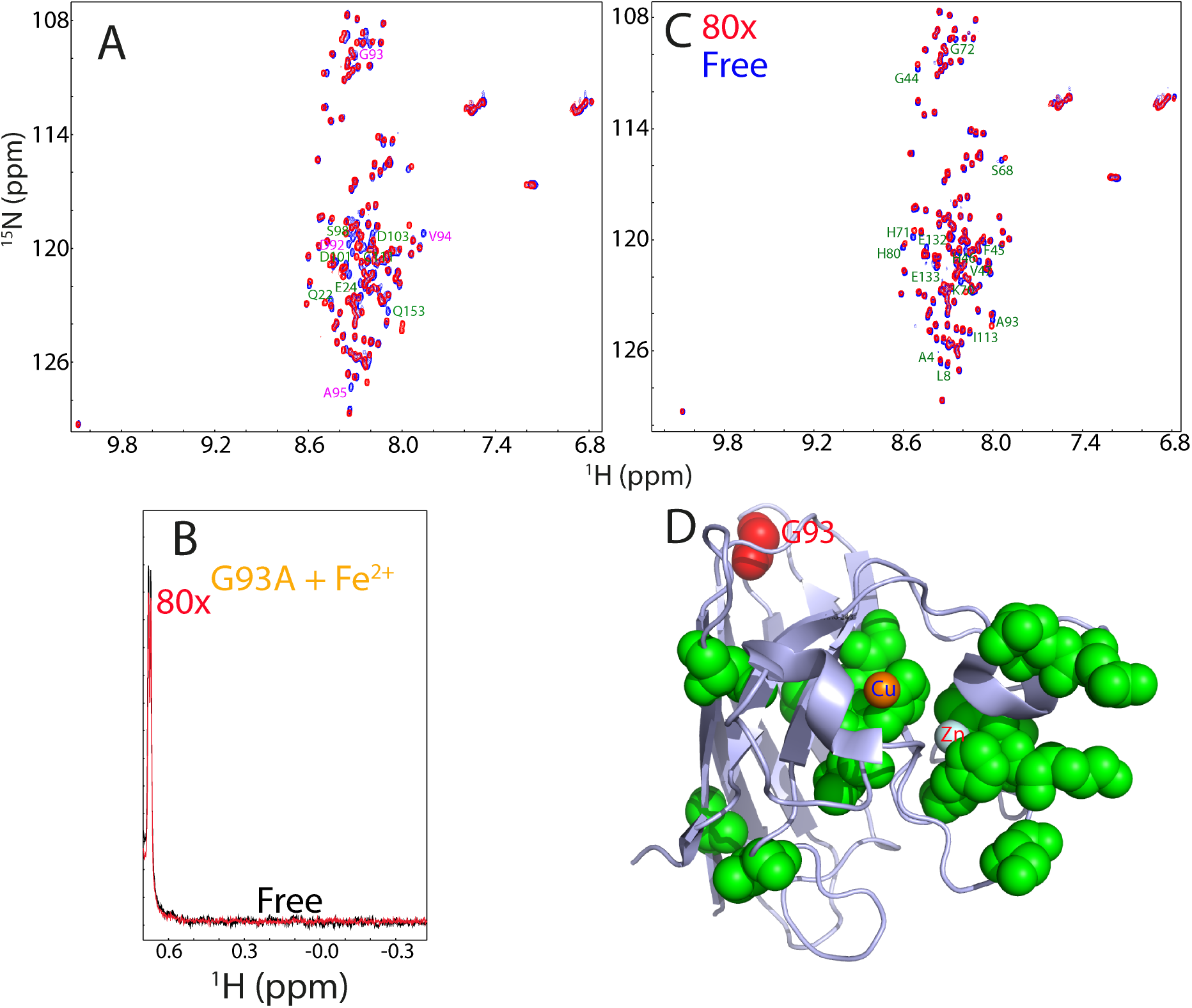
G93A-hSOD1 loses the folding capacity upon induction by Fe^2+^. (A) Superimposition of HSQC spectra of WT (blue) and G96A-hSOD1 (red). Residues with large shifts for their HSQC peaks are labeled. (B) Up-field NMR peaks (-0.5-0.64 ppm) of G93A-SOD in the absence (black) and in the presence of Fe^2+^ at a molar ratio of 1:80 (red). (C) Superimposition of HSQC spectra of G93A-hSOD1 in the absence (blue) and in the presence of Fe^2+^ at a molar ratio of 1:80 (red). Residues with significant shifts for their HSQC peaks are labeled. (D) The monomer structure of hSOD1 (1PU0) with the G93A-hSOD1 residues significantly perturbed by Fe^2+^ displayed in green spheres.

Amazingly, although the G93A mutation is not located within the Zn^2+^/Cu^2+^ binding pocket, G93A-hSOD1 lost the ability to undergo folding upon induction by Fe^2+^. Even in the presence of 80x Fe^2+^, no peaks characterized by the folded form manifested over up-field region (Fig. 8B), and in its HSQC spectrum (Fig. 8C). On the other hand, the presence of Fe^2+^ led to significant shifts of some HSQC peaks of the unfolded state, most of which are from the residues close to the Cu/Zn binding pocket except for Ala4, Leu6, Ile113 and the mutation residue Ala93 (Fig. 8C and 8C), indicating that Fe^2+^ might bind these residues of the unfolded state but initiate no further folding. However, again due to the paramagnetic nature of Fe^2+^, it is impossible to correlated these shifts to the solvent exposure, or/and direct binding, or/and binding-induced conformational/dynamic changes of these residues in the unfolded ensemble of G93A-hSOD1.

On the other hand, addition of Zn^2+^ is still able to trigger the formation of the folded state of G93A-hSOD1, as evident from the manifestation of up-field 1D (Fig. 9A) and well-dispersed HSQC peaks (Fig. 9B). However, as compared to the titration results of the WT hSOD1 by Zn^2+^, two significant differences were observed (Fig. 9A). First, at the same ratio, the intensity of up-field peaks of G93A is much lower than that of WT SOD1. Second, the Zn^2+^-induced formation of the G93A folded state will not become saturated even up to 1:60. However, attempts to further add Zn^2+^ triggered severe aggregation of G93A-hSOD1, thus preventing from further characterization at higher ratios of Zn^2+^. Nevertheless, both differences imply that the G93A-hSOD1 has a considerably reduced capacity in forming the folded form even induced by Zn^2+^.

**Fig. 9.**
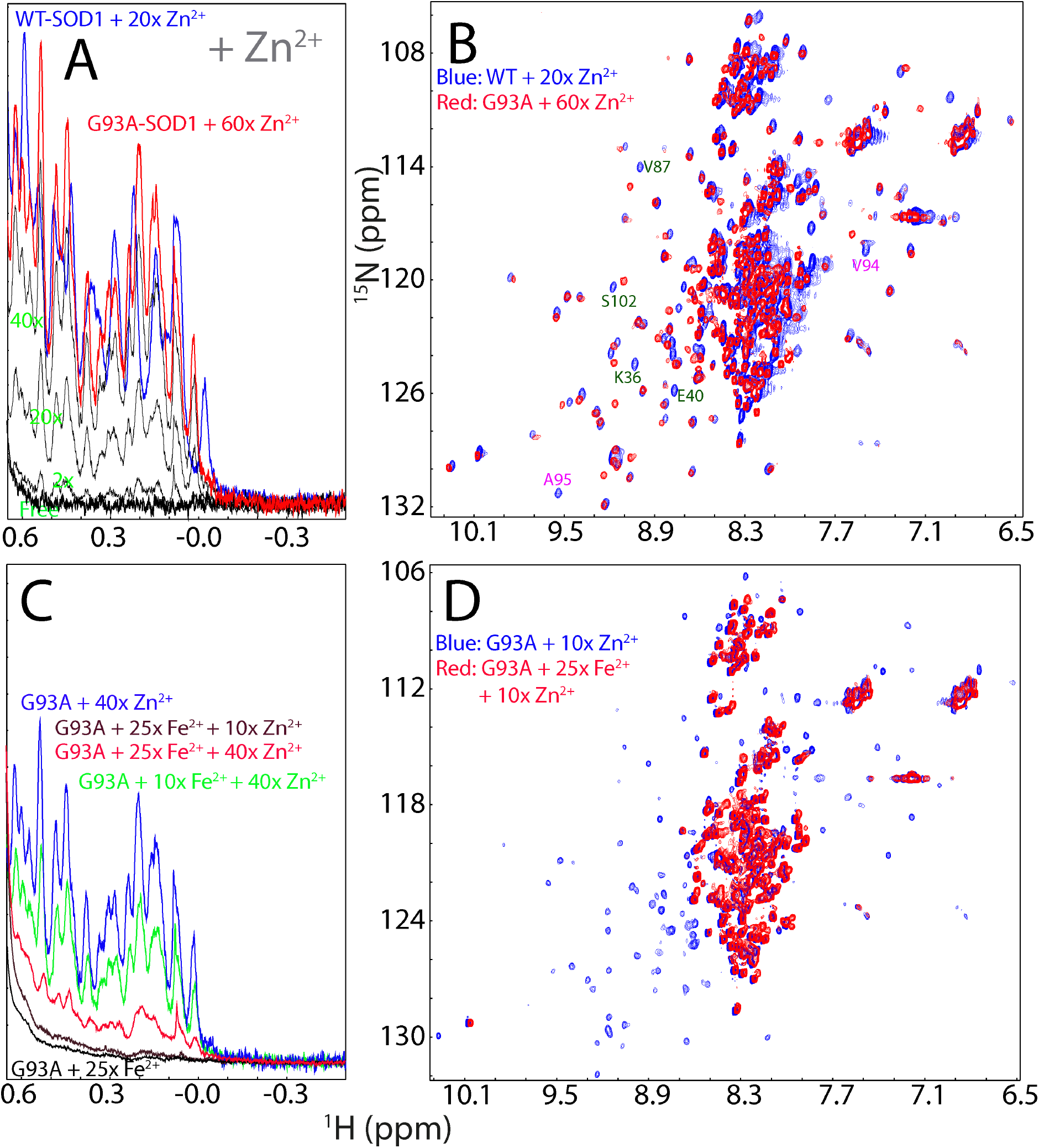
Fe^2+^ reduces the efficiency of the Zn2+-induced folding of G93A-hSOD1. (A) Up-field 1D NMR peaks characteristic of the folded WT or G93A-hSOD1 (-0.5-0.62 ppm) in the presence of Zn^2+^ at different molar ratios. (B) Superimposition of HSQC spectra of the WT hSOD1 in the presence of Zn^2+^ at a molar ration of 1:20 (blue), and G93A-hSOD1 in the presence of Zn^2+^ at a molar ration of 1:60 (red). Some residues with large shifts of their HSQC peaks are labeled (C) Up-field 1D NMR peaks characteristic of the folded G93A-hSOD1 (-0.5-0.62 ppm) with a pre-existence of 10x or 25x Fe^2+^, followed by addition of Zn^2+^ at different molar ratios. (D) Superimposition of HSQC spectra of G93A-hSOD1 in the presence of 10x Zn^2+^ without (blue) and with the pre-existence of 25x Fe^2+^ (red).

Human hSOD1 accounts for ∼1% of the total proteins in neurons. If assuming that the cellular protein concentration is ∼200 mg/ml, the hSOD1 concentration is 2 mg/ml, or 125 µM. On the other hand, it has been found that in the SOD1^G93A^ mice, the breakdown of blood–spinal cord barrier (BSCB) could result in the availability of free Fe^2+^ in neurons up to 800 ng/mg protein, which is ∼2.86 mM (22). That means that free Fe^2+^ concentration is ∼23 time higher than the SOD1 concentration in neurons. Therefore, here to mimic the in vivo situation, we evaluated the efficiency of Zn^2+^ in triggering the folding of G93A in the pre-existence of Fe^2+^ at ratios (G93A: Fe^2+^) of 1:10 and 1:25 respectively (Fig. 9C). Interestingly, even in the presence of only 10x Fe^2+^, the intensity of up-field peaks induced by adding 40x Zn^2+^ is much lower than that without Fe^2+^ (Fig. 9C). Most strikingly, in the pre-existence of 25x Fe^2+^, addition of 10x zinc is no longer able to induce up-field 1D (Fig. 9C), as well as well-dispersed HSQC peaks (Fig. 9D). Only upon addition of 40x zinc, the up-field peaks manifested to some degree, but however, those peaks are very weak and broad (Fig. 9C). Furthermore, visible aggregates were observed shortly in NMR tube and consequently, no high-quality HSQC spectrum could be acquired on this sample. Although the exact Zn^2+^ concentration in neurons remains to be defined, it was estimated that the Zn^2+^ concentration is only in the 10–30 μM range (38), which is not even reaching a ratio of 1 to 1 with hSOD1. As such, upon the breakdown of the blood–spinal cord barrier, because of the binding of Fe^2+^ to the unfolded G93A-hSOD1 ensemble, the maturation of the G93A-hSOD1 is expected to be significantly retarded in the neurons, consistent with the *in vivo* observations (5–8,19,20,22).

## Discussion

Despite exhaustive studies on hSOD1, here by a systematic assessment we have uncovered several previously-unknown results: firstly, in contrast to the widespread belief, the majority of 12 cations have no Zn^2+^-like ability to bind as well as to induce folding of nascent hSOD1. In particular, Ni^2+^, Cd^2+^ and Co^2+^, which have been widely believed to have physicochemical properties very similar to Zn^2+^ and thus utilized to substitute Zn^2+^ in mature hSOD1 for various biophysical and structural studies (5–7), have no detectable ability to bind as well as to induce the initial folding of nascent hSOD1. Intriguingly, although Cu^2+^, the other cofactor of hSOD1, shows extensive binding to the unfolded state of hSOD1. it only has weak capacity in inducing folding but owns a strong ability to trigger severe aggregation. This finding thus provides a potential mechanism for the extremely-critical role of Zn^2+^ in switching the toxic nascent hSOD1 associated with SALS into the non-toxic folded state, rationalizing the in-cell observation (39). Indeed, increasing evidence implies that abnormal zinc homeostasis is related to ALS pathogenesis and to increase the effective cellular concentration of Zn^2+^ such as by the dietary intake might offer an important therapy for ALS patients caused by hSOD1 (40).

Recently, we found that unexpectedly in addition to the positively-charged Zn^2+^, the negative-charged ATP and triphosphate could also induce folding of nascent hSOD1 at 1:8 ratio (24). By contrast, ADP, Adenosine, pyrophosphate and phosphate, as well as trimethylamine N-oxide (TMAO), the best-known inducer generally for protein folding, showed no detectable inducing capacity at concentrations where aggregation occurred for hSOD1 (24). Nevertheless, unlike the Zn^2+^-induced folded state, in the ATP-induced folded state, the loop regions around the Cu/Zn-binding pockets remain largely unformed (24). Therefore, Zn^2+^ appears to be selected by nature to serve as the cofactor of hSOD1 not just because it is a positively charged cation, but it owns a unique integration of at least three abilities: 1) to specifically coordinate the formation of the Zn^2+^-binding pocket and; 2) to induce folding of nascent hSOD1 as well as; 3) to occupy the pocket of mature hSOD1 to serve as a cofactor for enzymatic functions. In this context, while Ni^2+^, Cd^2+^ and Co^2+^ have the ability to bind the Zn^2+^-binding pocket in mature hSOD1, which needs the stabilization by the covalently cross-linked disulfide bridge Cys57-Cys146, they completely lack the Zn^2+^-like ability to induce folding of nascent hSOD1 and to specifically coordinate the formation of the Zn^2+^-binding pocket. On the other hand, although Cu^2+^ can site in a pocket in mature hSOD1 to serve as an enzymatic cofactor as well as extensively bind the unfolded state of nascent hSOD1, it has very weak capacity in inducing folding but exerts a strong adverse effect to trigger severe aggregation, which thus may represent one mechanism underlying the observation that the free Cu^2+^ ion is be highly toxic in cells and as such the human copper chaperone for hSOD1 (hCCS) is absolutely needed to carry and then load Cu^2+^ to hSOD1 (11–14).

Secondly, for the first time, we decode that out of 11 other cations, only Fe^2+^ has the Zn^2+^-like capacity to induce folding of nascent hSOD1. In particular, the results with WT, H80S/D83S and G93A mutants of hSOD1 reveal that although the Zn^2+^- and Fe^2+^-binding pockets are not identical, either binding by Zn^2+^ or Fe^2+^ is sufficient to induce folding of nascent hSOD1. This finding immediately raises an interesting question whether the Fe^2+^-bound hSOD1 exists or/and functions in any organisms. Indeed, there is iron superoxide dismutase (FeSOD), which was first discovered in *E. coli* in 1973 (41), shortly after the discovery of CuZnSOD in 1969 (42) and MnSOD in 1970 (43). Now FeSOD has been found in some bacteria, protists, mitochondria and chloroplasts but not in human. In fact, FeSOD is thought to be the first SOD to evolve due to the abundance of iron and low levels of oxygen in Earth’s primitive atmosphere. However, structurally FeSOD and MnSOD are almost identical but fundamentally different from CuZnSOD (44). As shown in Fig. S3, FeSOD (45) and MnSOD (46) have an identical two-domain structure rich in helices with Fe or Mn atom in the active site, while CuZn-SOD1 has a structure with a beta-barrel and a copper-zinc cluster in the active site. To the best of our knowledge, so far, there has been no report on detecting any FeSOD with the hSOD1 structural fold in any organisms. This raises a question of evolutionary interest: Why does nature engineer Cu/Zn-SOD1 which needs two cofactors and multiple steps to reach mature hSOD1 for functionality? In fact, to use Fe^2+^ as the SOD1 cofactor could potentially fulfill both requirements: initiating folding of nascent hSOD1 and catalyzing redox reactions in the mature enzyme.

Oxidative stress, resulting from an imbalance between the production of free oxidative radicals and the ability of the cell to remove them, has been increasingly identified to cause various human diseases, particularly neurodegenerative diseases (19–21). In this context, hSOD1 represents a central antioxidative enzyme while iron acts as a notorious accelerator. Iron is a double-edged sword: it is the most abundant “trace element” absolutely required for humans’ survival, but high levels of iron quickly lead to cell death. Under the normal conditions, the cellular concentration of free iron is very low, but under the pathological conditions such as the breakdown of the blood-central nerve system characteristic of neurodegenerative diseases and aging, the concentration of the blood-derived iron can reach very high (19–22,47–49). Indeed, iron is accumulated in various neurodegenerative as well as other diseases but the underlying mechanisms for its toxicity still remain to be fully elucidated. Currently, iron is known to trigger oxidative stress mainly through its reactivity with peroxide, thus generating the highly reactive hydroxyl radical by Fenton chemistry (49–56). Nevertheless, so far, there is no report even to imply that iron might manifest its cellular toxicity by specifically targeting SOD1.

Here, the competition experiments of Fe^2+^ and Zn^2+^ reveal that the presence of Fe^2+^ at high concentrations radically reduces the efficiency of the Zn^2+^-induced folding of both WT and in particular ALS-causing G93A hSOD1. Previously, the failure or even reduced efficiency of the folding has been proposed as a common mechanism for trapping the ALS-causing hSOD1 mutants in the highly-toxic species before the formation of the correct disulfide bridge, which are also prone to aggregation (13,18,31,35,36,38). Consequently, our finding implies a novel SOD1-dependent mechanism for the iron-induced production of oxidative stress by targeting the highly toxic mutants such as G93A, and even WT hSOD1 to disable its anti-oxidative functions.

In light of numerous previous results, we propose here a mechanism by which Fe^2+^ targets hSOD1 to provoke oxidative stress as well as to create toxic hSOD1 forms (Figure 10). More specifically, nascent hSOD1 exists as the unfolded state (Fig. 10A). Only upon supplement of Zn^2+^, a conformational equilibrium is established in which the folded population is formed (Fig. 10B), thus ready for further interacting with hCCS (Fig. 10C) to reach the mature and active hSOD1 (Fig. 10D) with copper loaded and the disulfide bridge formed capable of catalyzing the dismutation reaction (Fig. 10E). However, if in the presence of Fe^2+^ at high concentrations, even WT-hSOD1 associated with SALS may become the Fe^2+^-bound, which has an overall architecture similar to the Zn^2+^-bound (Fig. 10F and 10G), but unsuitable for further load of copper and formation of the disulfide bridge catalyzed by hCCS, As a consequence, the Fe^2+^-bound hSOD1 form without stabilization by the disulfide bridge would ultimately acquire high toxicity as found with other ALS-causing mutants including G93A (35) and H46R/H48Q (36), by becoming aggregated to form the iron-containing hSOD1 inclusion (Fig. 10H) which has been extensively observed in ALS patients.

**Fig. 10.**
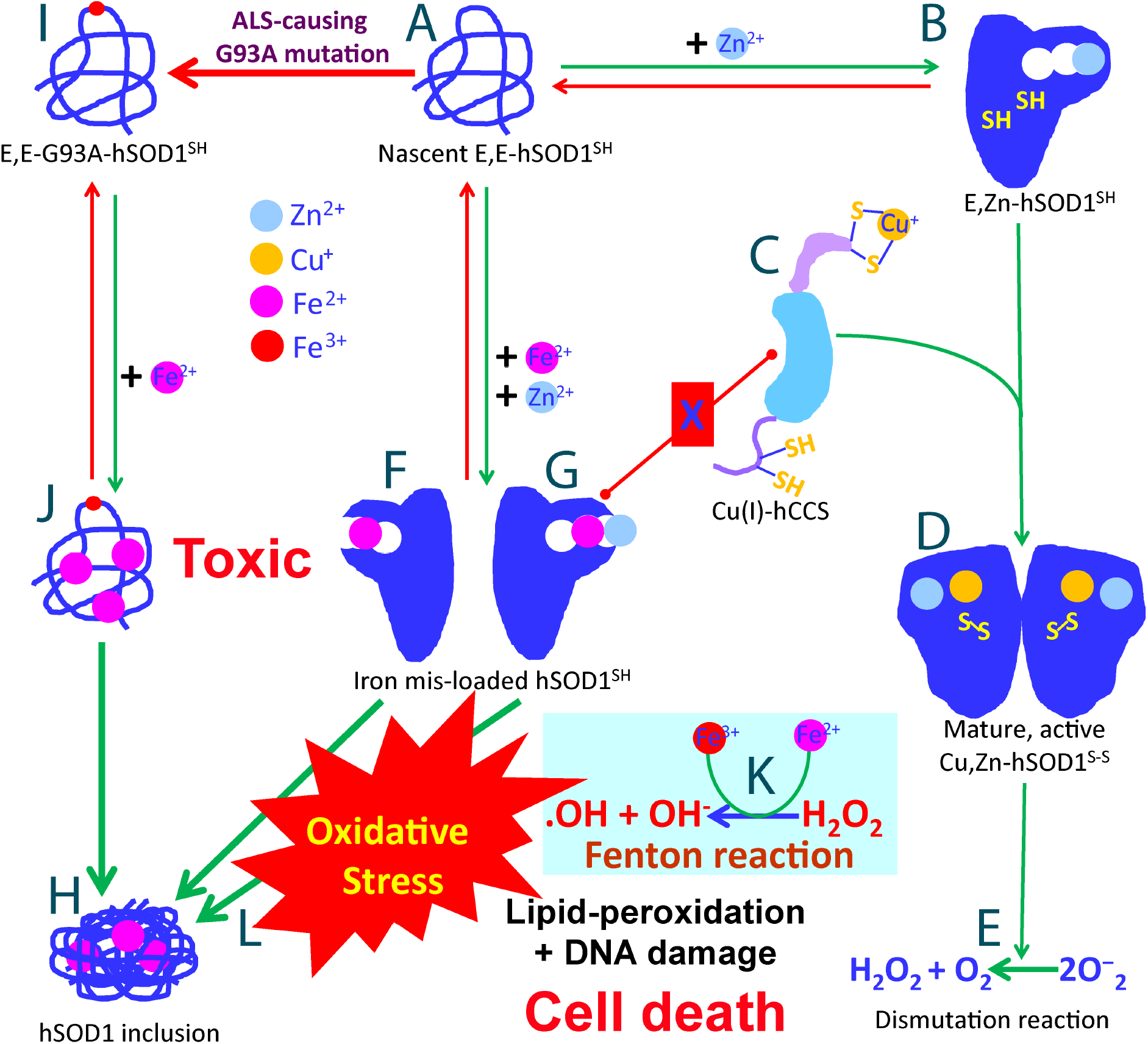
Speculative hSOD1-dependent mechanism by which Fe^2+^ provokes oxidative stress and traps both WT and ALS-causing mutant hSOD1 in the aggregation-prone and toxic forms.

Specifically, for G93A-hSOD1 causing FALS (Fig. 10I), even without the excess presence of Fe^2+^, the efficiency of the Zn^2+^-induced maturation has been demonstrated to be significantly reduced as previously reported (13,35) as well as observed here by NMR, likely owing to the reduced folding capacity induced by Zn^2+^ (Fig. 9A). This reduction is expected to provoke oxidative stress which may subsequently trigger the breakdown of blood–central nerve system barrier as previously demonstrated on the SOD1^G93A^ mice (47–56). Upon the breakdown, the concentration of the blood-derived Fe^2+^ could become as high as ∼25 times of the hSOD1 concentration. In this sense, although Fe^2+^ is no longer able to trigger the folding of G93A-SOD1, it can still bind to a set of residues of the unfolded G93A-hSOD1 ensemble (Fig. 10J) to dramatically disrupt the Zn^2+^-induced folding, thus blocking the initial folding critical for the G93A-hSOD1 maturation (Fig. 10J). As such, in the presence of Fe^2+^ at high concentration, the immature and disordered G93A-hSOD1 becomes accumulated for forming the iron-containing inclusion as observed *in vivo* (Fig. 10H). The iron-induced failure of the G93A maturation is expected to aggravate oxidative stress, which might further prompt the breakdown of blood– central nerve system barrier as a positive feedback loop. This mechanism also resolves a long-standing paradox that on the one hand, many mature ALS-causing mutants such as G93A-hSOD1 have their mature structures and activity almost identical to those of the mature WT hSOD1 (13,14,35), but on the other hand, *in vivo* they have been found to be highly toxic and aggregation-prone (13,14,48,52).

Our study also raises some interesting issues to be explored in the future. For example, it remains to investigate whether the Fe^2+^-bound hSOD1, which is chemically similar to Cu^+^-bound SOD1, also acquires the activity to catalyze endogenous production of nitric oxide to induce apoptosis, as previously detected on the zinc-deficient hSOD1 but with copper and disulfide bridge (35,36). Nevertheless, the residue-specific NMR results here unequivocally decipher that in addition to Fenton reaction (Fig. 11K), Fe^2+^ is also able to manifest its toxicity through a novel SOD1-dependent mechanism, by which Fe^2+^ traps the mutant or even WT hSOD1 into highly toxic forms to provoke significant oxidative stress (Fig. 10L). Both are expected to contribute to pathogenesis of neurodegenerative diseases including ALS as well as aging. This may partly rationalize the observation that under certain pathological conditions, the WT hSOD1 is also able to acquire cellular toxicity to initiate ALS. Furthermore, as hSOD1 exists in all human tissues, the SOD1-dependent mechanism we found here is expected to play general roles in triggering other diseases upon the bio-availability of Fe^2+^ at high concentrations in cells. Our study also provides mechanistic supports to the therapeutic approaches to treat neurodegenerative diseases or to slow down aging by repairing the breakdown of the blood-central nerve system barrier (56), or/and intaking iron-chelators by design from EDTA-like compounds or directly from antioxidant natural products such as flavonoids extensively existing in fruits and green tea, which can chelate and clean up Cu^2+^ and Fe^2+^ (57,58).

## Materials and methods

### Chemicals and preparation of the recombinant hSOD1 proteins

Chloride salts of 12 cations were all purchased from Sigma-Aldrich. The gene encoding the wild-type hSOD1 of the native sequence was purchased from Genscript with *E. coli* preferred codons (15,16). To remove the inference of extra residues, the gene was subsequently cloned into a modified vector pET28a without any tag (59). H80S/D83S and G93A mutants were generated by the use of QuikChange Site-Directed Mutagenesis Kit (Stratagene, La Jolla, CA, USA) (60). Subsequently the expression vector was transformed into and overexpressed in *Escherichia coli* BL21 (DE3) cells (Novagen). The recombinant WT and mutant hSOD1 proteins were all found in inclusion body. As a result, the pellets were first washed with buffers several times and then dissolved in a phosphate buffer (pH 8.5) containing 8 M urea and 100 mM dithiothreitol (DTT) to ensure complete conversion to Cys-SH. After 1 hour, the solution was acidified by adding 10% acetic acid and subsequently purified by reverse-phase (RP) HPLC on a C4 column eluted by water-acetonitrile solvent system (61–63). The HPLC elution containing pure recombinant hSOD1 was lyophilized and stored at -80 degree.

The generation of isotope-labeled proteins for NMR studies followed a similar procedure except that the bacteria were grown in M9 medium with the addition of (^15^NH_4_)_2_SO_4_ for ^15^N labeling (15,16). The purity of the recombinant proteins was checked by SDS–PAGE gels and verified by a Voyager STR matrix-assisted laser desorption ionization time-of-flight-mass spectrometer (Applied Biosystems), as well as NMR assignments. The concentration of protein samples was determined by the UV spectroscopic method in the presence of 8 M urea (15,16,64).

By exhaustively screening solution conditions including pH, types and concentrations of salts, we have successfully identified the optimized buffer to minimize the aggregation of nascent hSOD1 for high-resolution NMR studies: in 1 mM sodium acetate buffer at pH 4.5. This condition allowed the preparation of soluble and stable hSOD1 samples with a concentration up to 500 µM, which is required to collect high-quality triple-resonance NMR spectra. As hSOD1 of the native sequence contains four cysteine residues, previously we found that they started to form intermolecular disulfide bridges even in the presence of 10 mM DTT once the solution pH was above 5.0, and consequently resulted in immediate precipitation (15,16,65). Indeed, even for the C6A/C111S mutant with a significant reduction of the potential to form intermolecular disulfide bridges, its NMR sample still needed to be prepared at pH 5.0 (12). Here the low buffer-salt concentration (1 mM) was selected also to minimize the potential interference from the buffer cation (Na^+^) during titrations of 12 cations. Chloride salts of 12 cations were also dissolved in the same buffer with the final pH value adjusted to 4.5 by very diluted NaOH or HCl.

### ITC experiments

Isothermal titration calorimetry (ITC) experiments were performed using a Microcal VP isothermal titration calorimetry machine as described previously (40). The hSOD1 at a concentration of 5 µM was placed in a 1.8-ml sample cell, and each salt at a concentration of 6 mM were loaded into a 300-µl syringe. The samples were first degassed for 15 min to remove bubbles before the titrations were initiated. Control experiments with the same parameter settings were also performed for each inorganic salt without hSOD1, to subtract the effects because of sample dilution.

### NMR experiments

All NMR experiments were acquired at 25 °C on an 800 MHz Bruker Avance spectrometer equipped with pulse field gradient units as described previously (15,16). To achieve sequential assignments, triple resonance NMR spectra HN(CO)CACB and CBCA(CO)NH were collected on the ^15^N-/^13^C-double labeled hSOD1 samples while {^1^H}–^15^N heteronuclear steady-state NOE (hNOE) and HSQC-NOESY spectra were collected on the ^15^N-labeled samples. NMR ^1^H chemical shifts were referenced to external DSS at 0.0 ppm (15,16). NMR spectra were processed with NMR Pipe (66) and analyzed with NMR View (67).

For NMR titrations of 12 cations (68), ^15^N-labelled hSOD1 samples at a protein concentration of 50 μM in 1 mM sodium acetate-d3 buffer at pH 4.5 were used with stepwise additions of each cation at molar ratios of 1:0.5, 1:1, 1:2, 1:4, 1:6, 1:8, 1:10, 1:12, 1:16, 1:20, 1:24, 1:30 and 1:40 as well as higher ratios for mutant hSOD1 or Fe^2+^.

## Acknowledgements

This study is supported by Ministry of Education of Singapore MOE Tier 1 A-8000711-00-00 Grant to Jianxing Song.

## Author contributions

J.S. designed research; L. L., J. K. and J.S. performed research; L.L., J. K. and J.S. analyzed data; J.S. wrote the paper.

## Data availability statements

All data generated or analysed during this study are included in this published article and its supplementary information.

## Notes

### Competing Interest Statement

The authors have declared no competing interest.

### Summary of Updates

This version has included completely new results on two hSOD1 mutants H80S/D83S-hSOD1 and G93A-hSOD1, and reached new conclusions. The new results and conclusions have been presented in new Figures 2, 4, 7, 8, 9, 10.

## References

1. L. I. Bruijn, T. M. Miller, D. W. Cleveland, Unraveling the mechanisms involved in motor neuron degeneration in ALS. Annu. Rev. Neurosci. 27, 723–749 (2004).

2. M. C. Kiernan, et al. Amyotrophic lateral sclerosis. Lancet, 377, 942–955 (2011).

3. B. V. Zlokovic, Neurovascular pathways to neurodegeneration in Alzheimer’s disease and other disorders. Nat Rev Neurosci 12, 723–738 (2011).

4. D. R. Rosen, et al. Mutations in Cu/Zn superoxide dismutase gene are associated with familial amyotrophic lateral sclerosis. Nature. 362, 59–62 (1993).

5. Y. Sheng, et al. Superoxide dismutases and superoxide reductases. Chem Rev. 114, 3854–918 (2014).

6. J. S. Valentine, P. A. Doucette, S. Zittin Potter, Copper-zinc superoxide dismutase and amyotrophic lateral sclerosis. Annu Rev Biochem. 74, 563–93 (2005).

7. G. S. A. Wright, S. V, Antonyuk, S. S. Hasnain The biophysics of superoxide dismutase-1 and amyotrophic lateral sclerosis. Q Rev Biophys. 52, e12 (2019).

8. J. Huai, Z. Zhang, Structural Properties and Interaction Partners of Familial ALS-Associated SOD1 Mutants. Front Neurol. 10, 527 (2019)

9. O. Abel, et al. ALSoD: A user-friendly online bioinformatics tool for amyotrophic lateral sclerosis genetics. Hum Mutat. 33, 1345–1351 (2012).

10. F. Chiti, C. M. Dobson, Protein misfolding, functional amyloid, and human disease. Annu. Rev. Biochem. 75, 333–366 (2006).

11. L. Banci, et al. Human superoxide dismutase 1 (hSOD1) maturation through interaction with human copper chaperone for SOD1 (hCCS). Proc Natl Acad Sci U S A. 109, 13555–13560 (2012).

12. L. Banci, et al. Human SOD1 before harboring the catalytic metal: solution structure of copper-depleted, disulfide-reduced form. J Biol Chem. 281, 2333–2337 (2006).

13. L. Banci, et al. Atomic-resolution monitoring of protein maturation in live human cells by NMR. Nat Chem Biol. 9, 297–9 (2013).

14. E. Luchinat, et al. In-cell NMR reveals potential precursor of toxic species from SOD1 fALS mutants. Nat Commun. 5, 5502 (2014).

15. L. Lim, X. Lee, J. Song, Mechanism for transforming cytosolic SOD1 into integral membrane proteins of organelles by ALS-causing mutations. Biochim Biophys Acta. 1848, 1–7 (2015).

16. L. Lim, J. Song, SALS-linked WT-SOD1 adopts a highly similar helical conformation as FALS-causing L126Z-SOD1 in a membrane environment. Biochim Biophys Acta. 1858, 2223–30 (2016).

17. S. Sun et al. Translational profiling identifies a cascade of damage initiated in motor neurons and spreading to glia in mutant SOD1-mediated ALS. Proc Natl Acad Sci U S A. 112, E6993–7002 (2015).

18. A. Sannigrahi, et al. The metal cofactor zinc and interacting membranes modulate SOD1 conformation-aggregation landscape in an in vitro ALS model. Elife. 10, e61453 (2021)

19. Kwan JY, et al. Iron accumulation in deep cortical layers accounts for MRI signal abnormalities in ALS: Correlating 7 tesla MRI and pathology. PLoS ONE 7, e35241 (2012).

20. Ignjatović A, Stević Z, Lavrnić S, Daković M, Bačić G. Brain iron MRI: a biomarker for amyotrophic lateral sclerosis. J Magn Reson Imaging. 38, 1472–9 (2013).

21. Tokuda E, Furukawa Y. Copper Homeostasis as a Therapeutic Target in Amyotrophic Lateral Sclerosis with SOD1 Mutations. Int J Mol Sci. 2016 May; 17(5): 636.

22. Winkler EA, Sengillo JD, Sagare AP, Zhao Z, Ma Q, Zuniga E, Wang Y, Zhong Z, Sullivan JS, Griffin JH, Cleveland DW, Zlokovic BV. Blood-spinal cord barrier disruption contributes to early motor-neuron degeneration in ALS-model mice. Proc Natl Acad Sci U S A. 111, E1035–42 (2014).

23. C. G. Fraga, Relevance, essentiality and toxicity of trace elements in human health. Mol Aspects Med. 26, 235–44 (2005).

24. J. Kang, L. Lim, J. Song, ATP induces folding of ALS-causing C71G-hPFN1 and nascent hSOD1. Commun Chem. 2, 223 (2023).

25. Dyson HJ, Wright PE. Unfolded proteins and protein folding studied by NMR. Chem Rev 104(8):3607–3622 (2004).

26. R. W. Strange, et al. Variable metallation of human superoxide dismutase: atomic resolution crystal structures of Cu-Zn, Zn-Zn and as-isolated wild-type enzymes. J Mol Biol. 356, 1152–62 (2006).

27. Zhang O, Kay LE, Olivier JP, Forman-Kay JD (1994) Backbone 1H and 15N resonance assignments of the N-terminal SH3 domain of drk in folded and unfolded states using enhanced-sensitivity pulsed field gradient NMR techniques. J Biomol NMR 4(6): 845–858.

28. Qin H, Lim LZ, Song J. Dynamic principle for designing antagonistic/agonistic molecules for EphA4 receptor, the only known ALS modifier. ACS Chem Biol. 10,372–8 (2015).

29. Qin H, Lim LZ, Wei Y, Song J. TDP-43 N terminus encodes a novel ubiquitin-like fold and its unfolded form in equilibrium that can be shifted by binding to ssDNA. Proc Natl Acad Sci U S A. 111, 18619–24 (2014).

30. Palmer AG, Kroenke CD, Loria JP, Nuclear magnetic resonance methods for quantifying microsecond-to-millisecond motions in biological macromolecules. Methods Enzymol 339, 204– 238 (2001).

31. Roberts BR, Tainer JA, Getzoff ED, Malencik DA, Anderson SR, Bomben VC, Meyers KR, Karplus PA, Beckman JS. Structural characterization of zinc-deficient human superoxide dismutase and implications for ALS. J Mol Biol. 373, 877–90 (2007).

32. M. P. Williamson, Using chemical shift perturbation to characterise ligand binding. Prog Nucl Magn Reson Spectrosc. 73, 1–16 (2013).

33. J. Kang, L. Lim, J. Song, ATP binds and inhibits the neurodegeneration-associated fibrillization of the FUS RRM domain. Commun Biol. 2, 223 (2019).

34. Kayatekin C1, Zitzewitz JA, Matthews CR. Zinc binding modulates the entire folding free energy surface of human Cu,Zn superoxide dismutase. J Mol Biol. 384, 540–55 (2008).

35. Galaleldeen A, Strange RW, Whitson LJ, Antonyuk SV, Narayana N, Taylor AB, Schuermann JP, Holloway SP, Hasnain SS, Hart PJ. Structural and biophysical properties of metal-free pathogenic SOD1 mutants A4V and G93A. Arch Biochem Biophys. 492, 40–7 (2009).

36. Winkler DD, Schuermann JP, Cao X, Holloway SP, Borchelt DR, Carroll MC, Proescher JB, Culotta VC, Hart PJ. Structural and biophysical properties of the pathogenic SOD1 variant H46R/H48Q. Biochemistry. 48, 3436–47 (2009).

37. G. Otting, Protein NMR using paramagnetic ions. Annu Rev Biophys. 39, 387–405 (2010).

38. Frederickson CJ, Koh JY, Bush AI. The neurobiology of zinc in health and disease. Nat Rev Neurosci. 6, 449–62 (2005).

39. K. Homma, et al., SOD1 as a molecular switch for initiating the homeostatic ER stress response under zinc deficiency, Mol. Cell 52, 75–86 (2013).

40. H. F. Lopes da Silva, et al., Dietary intake and zinc status in amyotrophic lateral sclerosis patients. Nutr Hosp. 34, 1361–1367 (2017).

41. F. J., Jr. Yost, I. Fridovich. J. An Iron-containing Superoxide Dismutase from Escherichia coli Biol. Chem. 248, 4905–4908 (1973).

42. McCord, J. M.; Fridovich, I. Superoxide dismutase. An enzymic function for erythrocuprein (hemocuprein). J. Biol. Chem. 244, 6049–6055 (1969).

43. Keele, B. B., Jr.; McCord, J. M.; Fridovich, I. Superoxide dismutase from escherichia coli B. A new manganese-containing enzyme. J. Biol. Chem. 245, 6176–6181 (1970).

44. Stallings, W. C.; Pattridge, K. A.; Strong, R. K.; Ludwig, M. L. Manganese and iron superoxide dismutases are structural homologs. J. Biol. Chem. 259, 10695–10699 (1984).

45. Lah, M.S., Dixon, M.M., Pattridge, K.A., Stallings, W.C., Fee, J.A., Ludwig, M.L. Structure-function in Escherichia coli iron superoxide dismutase: comparisons with the manganese enzyme from Thermus thermophilus. Biochemistry 34: 1646–1660 (1995).

46. Borgstahl, G.E., Pokross, M., Chehab, R., Sekher, A., Snell, E.H. Cryo-trapping the six-coordinate, distorted-octahedral active site of manganese superoxide dismutase. J Mol Biol 296: 951–959 (2000).

47. Andrews NC, Schmidt PJ. Iron homeostasis. Annu Rev Physiol. 69, 69–85 (2007).

48. Montagne A, Barnes SR, Sweeney MD, Halliday MR, Sagare AP, Zhao Z, Toga AW, Jacobs RE, Liu CY, Amezcua L, Harrington MG, Chui HC, Law M, Zlokovic BV. Blood-brain barrier breakdown in the aging human hippocampus. Neuron. 85, 296–302 (2015).

49. B. Halliwell, J.M.C. Gutteridge, Free Radicals in Biology and Medicine, 3rd ed., Oxford Univ. Press, Oxford, (1999).

50. N. A. Simonian, J. T. Coyle, Oxidative stress in neurodegenerative diseases. Annu Rev Pharmacol Toxicol. 36, 83–106 (1996).

51. S. C. Barber, R. J. Mead, P. J. Shaw Oxidative stress in ALS: a mechanism of neurodegeneration and a therapeutic target. Biochim Biophys Acta. 1762, 1051–67 (2006).

52. E. A. Winkler, et al., Blood-spinal cord barrier disruption contributes to early motor-neuron degeneration in ALS-model mice. Proc Natl Acad Sci U S A. 111, E1035–42 (2014).

53. T. A. Rouault Iron metabolism in the CNS: implications for neurodegenerative diseases. Nat Rev Neurosci. 14, 551–64 (2013).

54. K. Joppe, et al. The Contribution of Iron to Protein Aggregation Disorders in the Central Nervous System. Front Neurosci. 13, 15 (2019).

55. A. Montagne, et al., Blood-brain barrier breakdown in the aging human hippocampus. Neuron. 85, 296–302 (2015).

56. P. T. Ronaldson, T. P. Davis. Targeting Transporters: Promoting Blood-brain barrier repair in Response to Oxidative Stress Injury. Brain Res. doi: 10.1016/j.brainres.2015.03.018 (2015).

57. X. Wang, et al. Role of Flavonoids in the Treatment of Iron Overload. Front Cell Dev Biol. 9, 685364 (2021).

58. E. Rodríguez-Arce, M. Saldías, Antioxidant properties of flavonoid metal complexes and their potential inclusion in the development of novel strategies for the treatment against neurodegenerative diseases. Biomed Pharmacother 143, 112236 (2021).

59. L. Wang, et al., TDP-43 NTD can be induced while CTD is significantly enhanced by ssDNA to undergo liquid-liquid phase separation. Biochem Biophys Res Commun. 499, 189–195 (2018).

60. Z. Wei, et al., Molecular mechanism underlying the thermal stability and pH-induced unfolding of CHABII. J. Mol. Biol. 348, 205–218 (2005).

61. M. Li et al., Nogo goes in the pure water: solution structure of Nogo-60 and design of the structured and buffer-soluble Nogo-54 for enhancing CNS regeneration. Protein Sci. 15, 1835–41 (2006).

62. M. Li, et al., Structural characterization of the human Nogo-A functional domains. Solution structure of Nogo-40, a Nogo-66 receptor antagonist enhancing injured spinal cord regeneration. Eur J Biochem. 271, 3512–22 (2004).

63. J Song, Tyrosine phosphorylation of the well packed ephrinB cytoplasmic beta-hairpin for reverse signaling. Structural consequences and binding properties. J Biol Chem. 278, 24714–24720 (2003).

64. C. N. Pace et al., How to measure and predict the molar absorption coefficient of a protein. Protein Sci. 4, 2411–2423 (1995)

65. J Song, et al., NMR solution structure of a two-disulfide derivative of charybdotoxin: structural evidence for conservation of scorpion toxin alpha/beta motif and its hydrophobic side chain packing. Biochemistry. 36, 3760–3766 (1997).

66. F. Delaglio et al., NMRPipe: a multidimensional spectral processing system based on UNIX pipes. J. Biomol. NMR. 6, 277–293 (1995).

67. B. A. Johnson, R. A. Blevins, NMR view: a computer program for the visualization and analysis of NMR data. J. Biomol. NMR. 4, 603–614 (1994).

68. Y. Liu, et al., Identification of a novel nonstructural protein, VP9, from white spot syndrome virus: its structure reveals a ferredoxin fold with specific metal binding sites. J Virol. 80, 10419–10427 (2006).

