## Supplementary Figures 1-3 for "Only Zn^2+^ and Fe^2+^ out of 12 cations can fold ALS-linked nascent hSOD1"

### Supplementary Materials

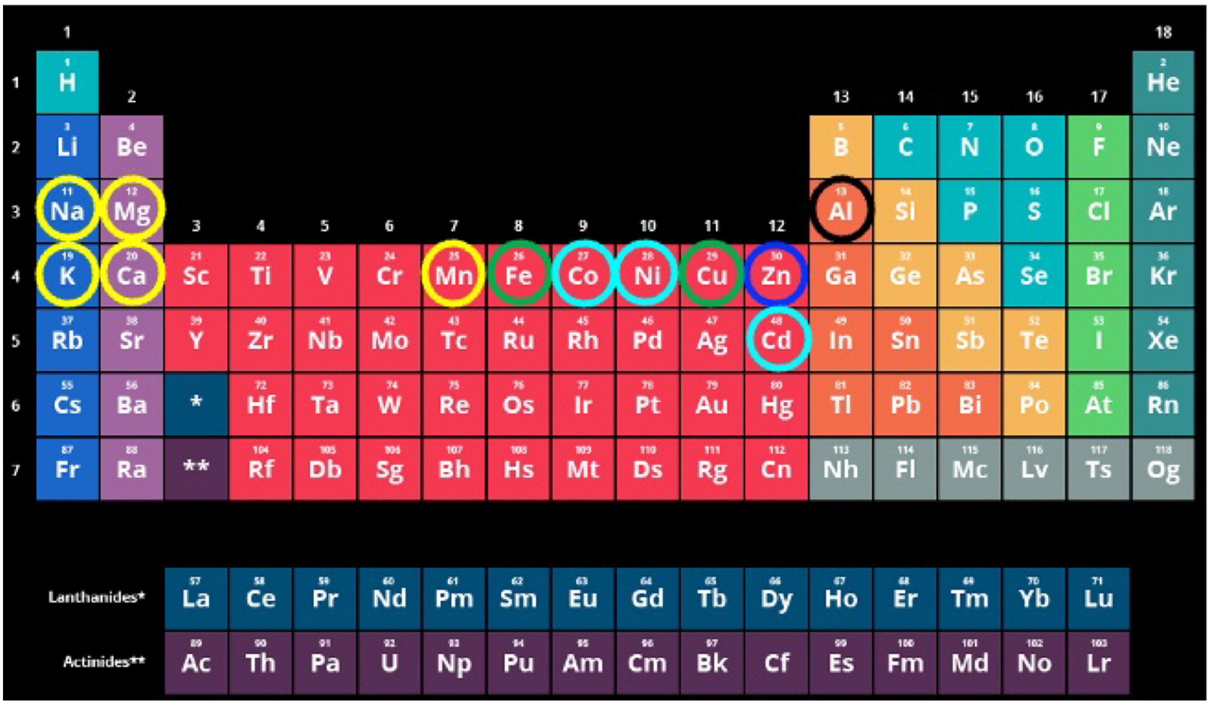


**Fig. S1. Selection of 12 cations.**
The periodic table of the elements to indicate the location of 12 metal cations selected for the present study.

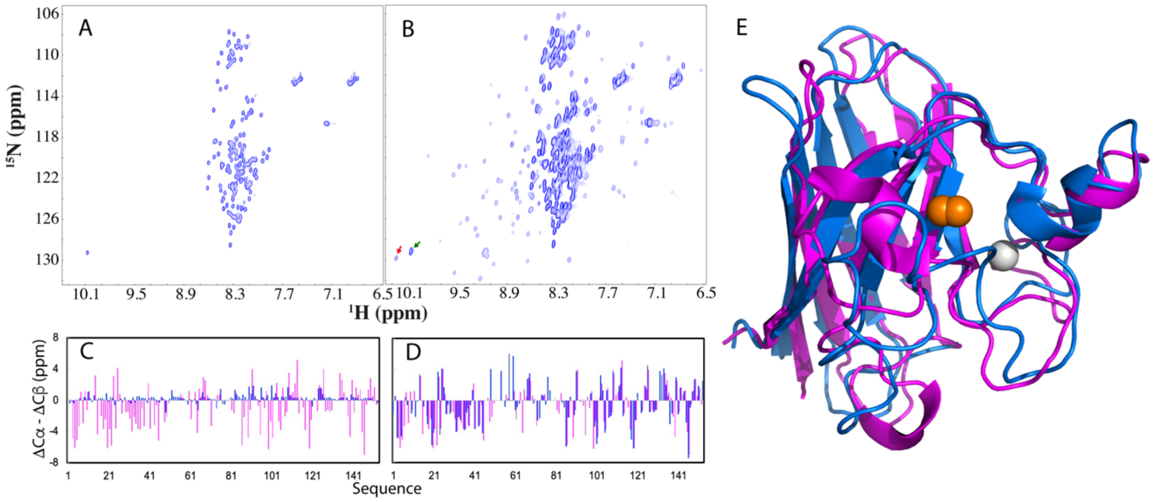


**Fig. S2. $Zn^{2+}$-induced folding of nascent hSOD1.**
HSQC spectra of nascent hSOD1 in the absence (A) and in the presence of $Zn^{2+}$ at a molar ratio of 1:20 (B). (C) Residue specific (ΔCα–ΔCβ) chemical shifts of nascent hSOD1 (blue) and in the presence of $Zn^{2+}$ at a molar ratio of 1:20 (purple). (D) Residue specific (ΔCα–ΔCβ) chemical shifts of the $Zn^{2+}$-induced hSOD1 and those of C6A/C111S (blue) (BMRB Entry of 6821). (E) Superimposition of the NMR structure of the super-stable pseudo-WT hSOD1 C6A/C111S (PDB ID of 2AF2) without the disulfide bridge and copper (blue) and crystal structure (PDB ID of 2C9V) of mature hSOD1 (purple). Zinc ion is in grey sphere and copper in orange sphere.

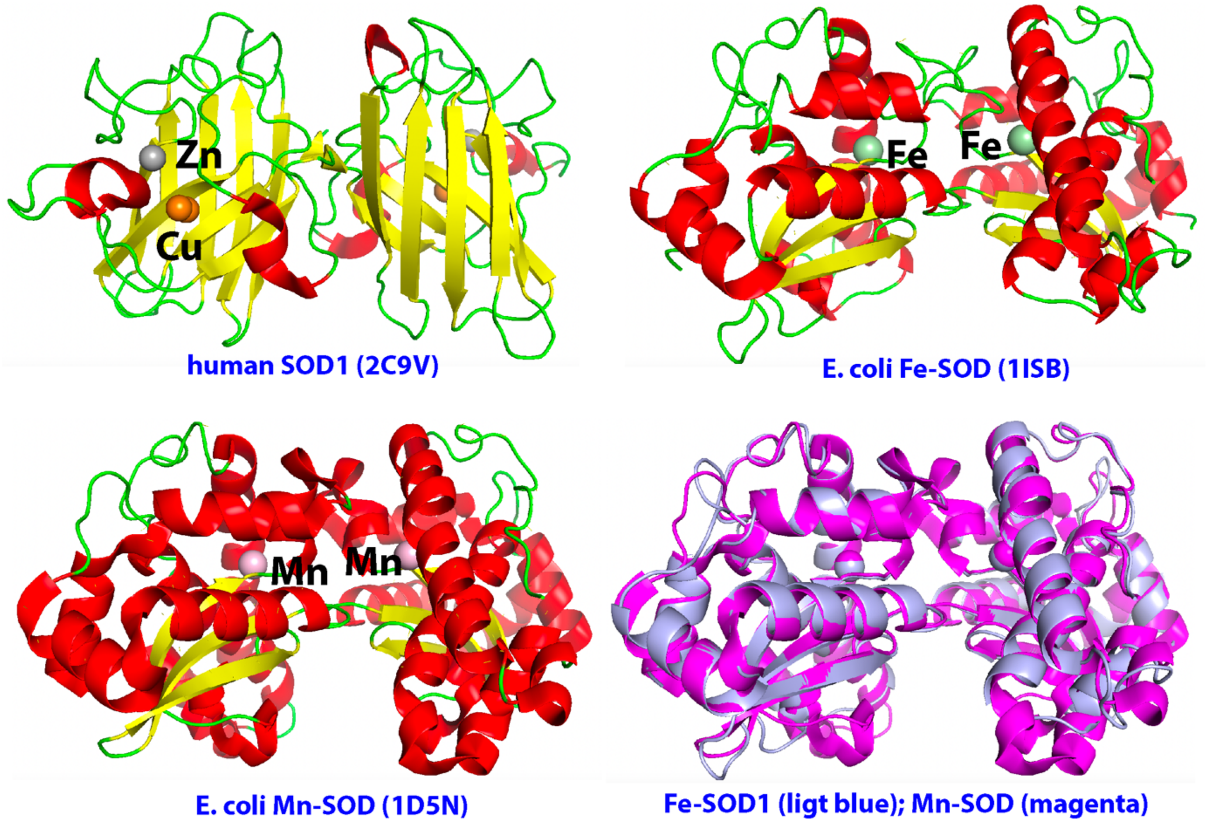


**Fig. S3. Three-dimensional structures of human CuZn-superoxide dismutase 1 (hSOD1), as well as *E. coli* Fe-SOD and Mn-SOD.**
